# Compound Mechanism of Action and Polypharmacology can be Elucidated by Large-Scale Perturbational Profile Analysis

**DOI:** 10.1101/2023.10.08.561457

**Authors:** Lucas ZhongMing Hu, Tal Hirschhorn, Stanley I. Goldstein, Diana Murray, Stephen J. Trudeau, Mikko Turunen, Eugene Douglass, Sergey Pampou, Adina Grunn, Léo Dupire, Ronald Realubit, Albert A. Antolin, Alexander L.E. Wang, Hai Li, Prem Subramaniam, Prabhjot S Mundi, Charles Karan, Mariano Alvarez, Andrew Emili, Barry Honig, Brent R. Stockwell, Andrea Califano

## Abstract

Drug Mechanism of Action (MoA) is often represented as a small repertoire of tissue-independent, high-affinity binding targets. Yet drug activity and polypharmacology are also manifested through a broad range of off-target and tissue-specific effector proteins. To address this dichotomy, we generated a compendium of drug perturbation profiles for >700 oncology drugs in cell lines representing high-fidelity models of 23 aggressive tumor subtypes, as well as an integrative framework for the proteome-wide assessment of drug-mediated, tissue-specific differential protein activity. Extending MoA and polypharmacology assessment via proteome-wide, multi-tissue analyses revealed novel polypharmacology, including for MAPK, PI3K, and folate pathways inhibitors—as supported by Western blots, thermal shift assays, and structural analysis—as well as functional inhibitors of challenging oncoprotein targets, including MYC, CTNNB1, CENPF and UHRF1. The dataset and associated computational framework might be extensible to additional compound libraries, supporting global, quantitative approaches to defining the mechanisms of action of many drugs.

## Introduction

Targeted therapy and precision medicine are predicated on the ability to accurately elucidate the mechanisms of action (MoA) of small molecule compounds and to match them to disease-specific dependencies. However, the binding affinity and molecular structures of targets revealed by biochemical assays (*on-target*) do not provide comprehensive, proteome-wide characterization. Moreover, drug efficacy and toxicity are increasingly linked to complex polypharmacology effects—mediated by additional low-affinity binding proteins (*off-targets*), tissue-specific indirect effector proteins (*secondary targets*), and safety-related proteins (anti-targets)—the detection of which by traditional assays is challenging^1^. Indeed, MoA studies, including those based on recent advances in pull-down technology^2^, often rely on *in vitro* affinity binding assays designed to investigate only specific protein classes, such as kinases^3^, failing to reveal a compound’s full range of high-affinity and low-affinity binding proteins and, perhaps more importantly, tissue-specific effectors, including those implicated by cell-type-specific canalization mechanisms; such polypharmacology and downstream effects can be thought of as a broader field effect of a drug rather than its one mechanistic high affinity binding protein. To avoid confusion with the canonical MoA concept, we will refer to this additional level of characterization as “*extended MoA”* (eMoA). Indeed, elucidating the proteome-wide, context-specific “footprint” of a drug, which ultimately determines its potential clinical efficacy and toxicity, has emerged as a critically-relevant, yet still elusive endeavor in precision medicine^3,4^, including in oncology-related applications^1,5,6^.

To address this challenge, we generated a large-scale repertoire of genome-wide, tissue-specific perturbational profiles and analyzed them using a framework built upon VIPER, an extensively validated algorithm for differential protein activity assessment^7–12^. The resulting Pan-Cancer Assessment of Compound Effectors and Activity (PanACEA) compendium and analytical framework leverage >18,000 RNA-seq profiles and ∼50 million drug-mediated, differential protein activity measurements, thus representing the largest genome and proteome-wide resource of its kind. Attesting to the quality and informative nature of these data and associated analytical framework, VIPER-based protein activity profiles were instrumental in identifying drugs targeting tumor-specific dependencies in multiple pre-clinical studies, across a wide range of aggressive human malignancies^9,10^, as well as in a clinical context^13^.

Perturbational profiles, characterizing the response of a tissue to one or more small molecules, have been previously leveraged to assess compound MoA. These include proteomic assays^14–17^—for instance, to elucidate signal-transduction protein targets^18^—and transcriptional profiles focusing on changes in transcriptional cell state following drug perturbation^5,6,19^. Among these, proteomic assays are still expensive, labor intensive, and sparse. As such, they have been restricted to study only a handful of compounds of interest in specific cellular contexts, rather than large compound libraries^14,15^, or have focused only on specific cell lines^16^. In addition, these assays report on protein abundance, which represents only an indirect measure of a protein’s enzymatic or regulatory activity. Transcriptional profiles are less expensive, more scalable, and more amenable to AI and automation. However, their ability to directly inform drug-mediated changes in protein activity or abundance has been limited. In addition, their use has been restricted either to small, landmark gene sets^19^ or a handful of tissues^6,20^.

To transform genome-wide RNA-seq profiles from drug perturbations into accurate protein activity profiles that optimally inform on drug eMoA, we leveraged the VIPER algorithm^7^. Akin to a highly multiplexed gene reporter assay, VIPER was designed to measure each protein’s differential activity, based on the differential expression of its tissue-specific targets (*regulon*). VIPER protein activity measurements compare favorably to antibody-based measurements^12,21^ and can be used to effectively monitor the activity of >6,500 regulatory and signaling proteins, including from single-cell profiles. The experimental component of this study is based on the high-throughput generation of comprehensive RNA-seq profiles, representing the response of 23 cell lines to a library of more than 700 clinically relevant drugs. VIPER was then used to transform gene expression profiles into proteomic profiles representing the differential activity of transcription factors and co-factors—henceforth transcriptional regulators (TRs)—and signaling proteins, in drug-treated vs. DMSO-treated samples.

While large-scale perturbational profile libraries, such as LINCS^19^ and Tahoe^22^, are available, they have limitations for the protein-level elucidation of drug eMoA, including suitability to VIPER-based analyses. Specifically, (a) they only report the effect on a small set (∼1,000) of landmark genes (LINCS) or use low-depth (∼10000 reads/cell) single cells profiles (Tahoe); (b) perturbations were performed at isomolar concentrations (*e.g.*, 1 μM, 5 μM, etc.), thus potentially missing relevant pharmacological windows where compound activity is optimally manifested; (c) perturbed cell lines were not carefully selected as high-fidelity models of human disease; (d) compound libraries contain few clinically-relevant drugs, such as FDA-approved and late-stage developmental compounds (*i.e.*, in phase 2 or 3 clinical trials); and (e) cell lines were co-cultured, thus affecting mechanisms via paracrine interactions (Tahoe). Thus, the resulting reduced-representation profiles cannot be effectively converted to protein activity profiles using VIPER because only 5% of each protein’s targets are available, on average.

In contrast, we leveraged PLATE-Seq—a low-cost, fully-automated, genome-wide RNA-seq profiling technology^23^, thus supporting VIPER analyses that are virtually indistinguishable from those based on high-depth TruSeq profiles^7^. In addition, the 23 cell lines comprising this initial PanACEA release—as well as additional cell lines being added to this resource—were carefully selected as high-fidelity models of human cancer subtypes, based on the overlap of their Master Regulator (MR) proteins with those already shown to be ultra-conserved in specific tumors subtypes^24^ (using the OncoMatch algorithm^25,26^). This is critical because we have shown that the eMoA of drugs—as predicted from perturbational profiles from high-fidelity models *in vitro*—is faithfully recapitulated in PDX models *in vivo*^10^. Finally, as proposed^6,27^, this design titrates compounds at their highest sublethal concentration (48 h EC_20_)—as assessed from systematic drug response curves (DRC)—thus avoiding concentrations that are too high—thus confounding the analysis by engaging cell stress and death pathways—or too low to reveal mechanism.

PanACEA profiles are thus expected to effectively support *de novo*, tissue-specific drug eMoA characterization, including: (a) cell-context-specific canalization of different drug perturbations, resulting in activation of common response mechanisms, (b) proteome-wide assessment of drug eMoA similarity, (c) elucidation of drug polypharmacology, and (d) discovery of novel inhibitors of undruggable proteins, including those previously identified as critical dependencies of molecularly distinct tumor subtypes^24^.

The resulting integrative analysis framework identified tissue-context-specific inhibitors of challenging targets, such as CTNNB1, UHRF1, MYC and CENPF, previously identified as tumor-specific dependencies^28^. It also allows prediction and validation of both compound polypharmacology and eMoA of poorly characterized compounds. For example, as predicted by the analysis, we confirmed that (a) Kw2499, originally characterized as a FLT3/ABL/Aurora kinase inhibitor, also potently inhibits the PIK3CA/AKT pathway, (b) biib021 (an HSP90 inhibitor^29^) and Az12419304 (a compound with unknown activity) represent effective MEK pathway inhibitors, and (c) Mgcd265, a multi-kinase inhibitor, plays a critical role targeting folate metabolism. To further assess the mechanisms that support these functional effects, we used mass-spectrometry-based Proteome Integral Solubility Alteration (PISA)^30^, the PrePCI database of protein-compound intreactions^31,32^, and Induced Fit Docking (IFD) analyses^33^ (see STAR Methods). Taken together, these revealed that Kw2449 stabilizes RPS6KB1, a core protein in PIK3CA/AKT pathway, and Az12419304 inhibits the MEK pathway through direct engagement with BRAF. We also found that Mgcd265 destabilizes the GP-transamidase (GPIT) complex, a well-known folate receptor (FR) anchor^34^, to affect cellular folate uptake, which was validated by a folate uptake assay.

The proposed approach is fully generalizable and can be adapted to studying the eMoA of novel chemical libraries in other physiologic or pathologic contexts, from neurological and neurodegenerative diseases^35^ to targeting mechanisms associated with cell state reprograming^36^ and infections^37^. The PanACEA resource, including both RNA-seq and protein activity profiles, is freely available to the research community.

## Results

### PanACEA Database Assembly

PanACEA comprises n = 18,046 RNA-Seq profiles representing the genome-wide transcriptional response of 23 cell lines 24 h following treatment with >700 FDA-approved and late-stage investigational compounds, as well as vehicle control media, in duplicate. The profiling protocol was specifically designed to optimize for elucidation of tissue-specific drug “extended Mechanism of Action” (eMoA) and polypharmacology; see **Figure 1** for an illustrative workflow of the study methodology.

**Figure 1:**
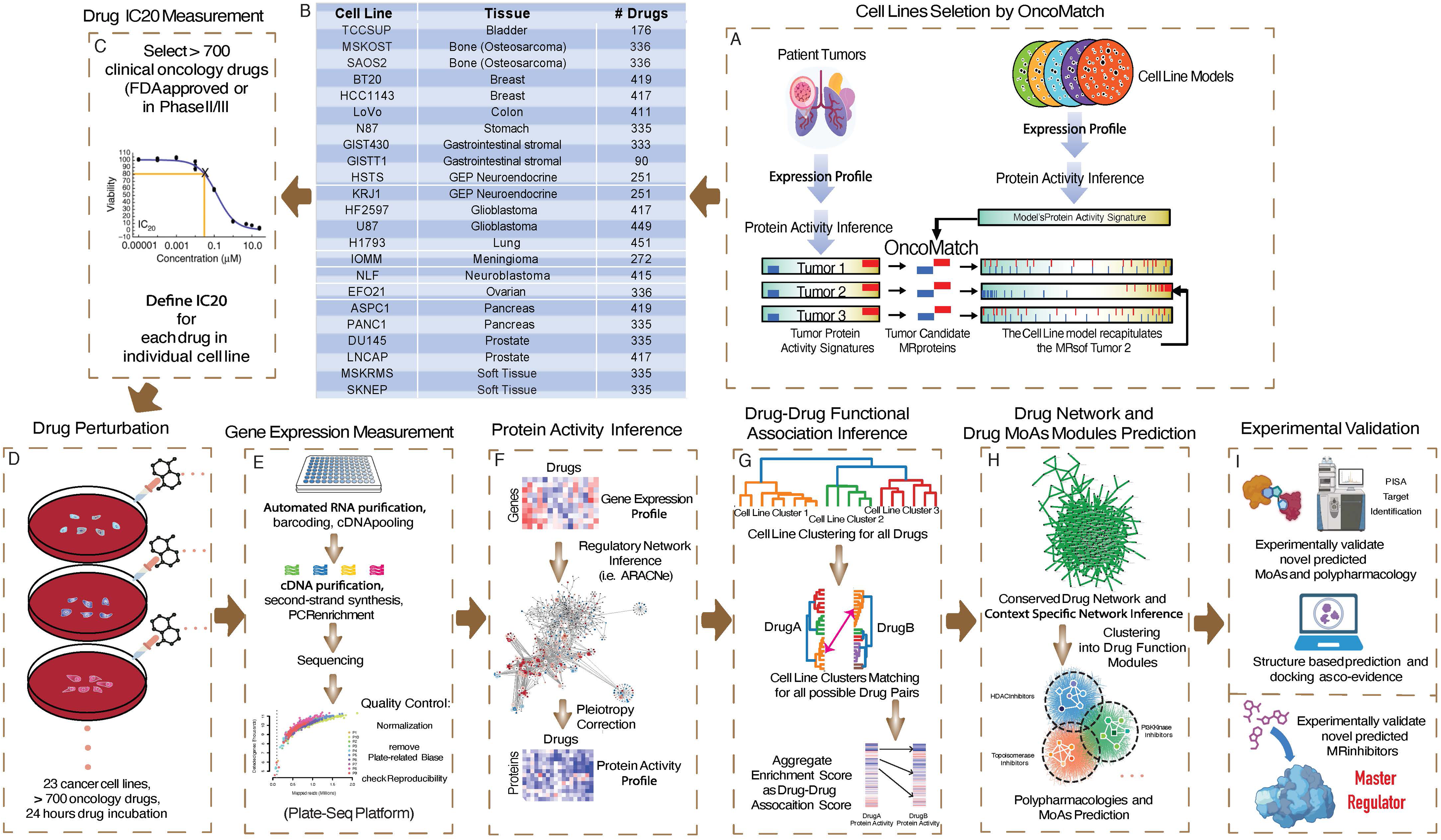
Schematic view of the PanACEA resource for the characterization of oncology drug eMoA. (A) The OncoMatch algorithm was used to select CCLE cell lines that recapitulate the MR signature of human tumor cohorts. (B) Compendium of cell lines, associated tissue of origin and number of drugs they were treated with to produce the PanACEA resource. (C) The EC_20_ of each drug in each cell line was assessed from 10-point dose response curves. (D) 23 selected cancer cell lines were treated with >700 distinct oncology drugs (in total) for 24 hours at their established EC_20_. (E) PLATE-Seq was used to generate genome-wide RNA-Seq profiles following each drug perturbation. (F) The ARACNe and VIPER algorithms were used to compute differential protein activity in drug vs. vehicle control-treated cells. (G) Drug-drug eMoA similarity was assessed and clustered to generate conserved activity modules. (H) A functional Drug eMoA Similarity Network was generated to visualize same-MoA drug modules suggesting polypharmacology or novel activity prediction. (I) Predicted eMoA, polypharmacology, and MR-specific activity was validated for selected drugs.

Critical differences between PanACEA and Connectivity Map (CMap)^20^, LINCS (L1000)^19^, and Tahoe^22^ profiles include: (a) PanACEA comprises both genome-wide RNA-Seq profiles and VIPER-assessed differential protein activity profiles for ∼6,500 regulatory and signaling proteins; (b) PanACEA was designed to support clinical translation by (*i*) selecting cell-lines representing high-fidelity models of rare and/or aggressive malignancy subtypes (**Figure 1A-B**), and (*ii*) including a vast majority of clinically-relevant oncology drugs—including FDA-approved and late-stage experimental agents in Phase 2 and 3 clinical trials (**Table S1**); and (c) drug-concentrations were chosen as the 48 h, cell-line specific EC_20_, as assessed from 10-point dose-response curves (DRC) (**Figure 1C**). While the latter adds significant complexity and cost to the study design, it is also critical to avoid confounding effects associated to cell stress/death pathway activation—accounting for up to 75% of the variance in existing drug-perturbation datasets. This confounds eMoA elucidation by masking critical targets, while artificially increasing drug eMoA similarity^38,39^.

The key advantage of this design is its optimal suitability to VIPER-based analyses, aimed at evaluating drug-mediated changes in transcriptional regulator (TR) and signaling protein activity^7^. Notably, of the 23 cell lines in this study, only two (LoVo and BT20) are included in LINCS^19^. This is important because PanACEA cell lines were selected to represent human tumors, based on the analysis of either published databases or individual patients enrolled in the N of 1 study at Columbia, representing rare, aggressive adult/pediatric malignancies^10^.

### High-fidelity cell line model selection

To support their ability to survive *ex vivo*, cell lines accumulate genetic, epigenetic, and aneuploidy artifacts that are absent in their parental tumors. While poorly characterized and understood, these events can significantly affect the pathophysiologic fidelity of the model, including their response to drugs. As a result, selecting cell lines for perturbational analysis based on the either (a) the presence of specific driver mutations, (b) the activity of specific surface markers, (c) *in vitro* growth kinetics, or (d) morphology criteria may be suboptimal. For instance, the commonly used strategy of selecting cell lines with good growth kinetics can bias their sensitivity to cell-cycle-dependent chemotoxic agents, thus masking subtler yet more critical dependencies in slower-growing human tumors.

As previously shown^24^, VIPER-inferred MRs representing mechanistic determinants of cancer cell state are ultra-conserved among tumors representing the same subtypes; in contrast, they are often not recapitulated by their cognate cell lines. For instance, among several neuroblastoma cell lines harboring a *MYCN* amplification, only three were shown to effectively recapitulate the MR protein panel hyper-conserved in *MYCN*-amplified human tumors. These cells were instrumental in recapitulating TEAD4 essentiality *in vivo*^11^.

Based on these observations, we developed^11,25^ and validated^9,10,40^ the OncoMatch algorithm to assess *model fidelity* based on the statistical significance of MR conservation in a model/tumor pair. OncoMatch tests for the enrichment of the top 50 MRs—*i.e.*, 25 most activated (MR↑) and 25 most inactivated (MR↓)—in a patient tumor in proteins differentially active in a cancer model (*e.g.*, cell line, PDX, organoid, etc.) and vice-versa. We have shown that this metric effectively helps predict drug effects on protein activity that are recapitulated *in vivo*^10^. Moreover, it is highly stable when the number of selected MRs varies between 10 and 200, see^10^.

We used OncoMatch to optimize selection of high-fidelity cell lines for perturbational studies, thus making them more relevant to clinical translation (STAR Methods). The 23 PanACEA cell lines include: H1793 (lung adenocarcinoma, LUAD), LNCAP and DU145 (AR-dependent and AR-independent prostate adenocarcinoma, PRAD), PANC1 and ASPC1 (morphogenic and gastrointestinal lineage states of pancreatic ductal adenocarcinoma, PAAD), EFO21 (ovarian serous cystadenocarcinoma, OV), TCCSUP (bladder urothelial carcinoma, BLCA), HCC1143 and BT20 (basal-like breast cancer, TNBC), N87 (stomach adenocarcinoma, STAD), LoVo (colon adenocarcinoma, COAD), HSTS and KRJ1 (gastroenteropancreatic neuroendocrine tumor, GEP-NET), IOMM (meningioma, MNG), MSKOST and SAOS2 (osteosarcoma, OS), SKNEP (Ewing sarcoma, EWS), MSKRMS (rhabdomyosarcoma, RMS), NLF (neuroblastoma, NB), GISTT1 and GIST430 (imatinib-sensitive and resistant gastrointestinal stromal tumor, GIST) and HF2597 and U87 (proneural and mesenchymal glioblastoma, GBM); see STAR Methods.

### Perturbational profile generation

Cells were first cultured and expanded using the recommended culture media (STAR Methods), until enough were available to seed 96 or 384-well plates. 10-point Dose Response Curve were then generated for drug libraries comprising 90 to 451 compounds, depending on cell type (**Table S1**). While the number of drugs/cell line is variable—since profile generation has been ongoing for the last 10 years—most cell lines where treated with ∼140 FDA approved antineoplastics, ∼170 late-stage experimental drugs, and a variable number of tool compounds from the Tocris and MicroSource collections with EC_20_ ≤ 2μM—as required for optimal utilization of 96- or 384-well PLATE-Seq plates—as well as vehicle control media (DMSO).

Additional chemonaïve cells were then expanded, plated again, at an average of 4,000 cells per well, and allowed to rest for 24 h before treatment with the cell line-specific 48 h EC_20_ of each compound. Cells were then harvested at 24 h following treatment, enzymatically processed for library preparation, using the PLATE-Seq methodology, and sequenced to a depth of 1M to 2M reads; RNA-seq profiles then underwent quality control and normalization (STAR methods).

### Data Reproducibility

To assess gene expression profile reproducibility, we leveraged the LoVo and BT20 cell lines, which are also included in LINCS. As shown in **Figure S1 A-D**, the differences in Pearson and Spearman correlation between technical replicates (*i.e.*, same cell line and drug), as assessed for the same set of ∼1,000 LINCS landmark genes, were dramatically higher in PanACEA compared to LINCS, as assessed by the conservative, non-parametric Wilcoxon test. For LoVo, Wilcoxon p-values were p = 2.75×10^-111^ (Pearson) and p = 6.77×10^-47^ (Spearman), while, for BT20, they were p = 4.71×10^-186^ (Pearson) and p = 4.39×10^-128^ (Spearman). A more critical point is that PanACEA comprises genome-wide expression profiles, while LINCS is limited to ∼1,000 landmark genes, thus relying on computational inference for the others. This is potentially problematic since inference methods were trained on steady state populations rather than on those undergoing complex dynamics following perturbation.

Next, as suggested by recent publications^24^, we asked whether VIPER analyses could further improve reproducibility. This could not be done for LINCS profiles, since use of landmark genes decreases regulon coverage by ∼20-fold, thus preventing VIPER analyses. TRs and signaling protein regulons were inferred by analyzing RNA-seq profiles from histology-matched human cohorts using ARACNe^41^ (see **Table S2** for the corresponding tumor cohort list). Reproducibility was assessed by GSEA analysis of differentially expressed genes and differentially active proteins, see STAR Methods. Protein activity was significantly more reproducible than gene expression, in all but one cell line, with improvement statistics ranging from p = 1.13×10^-13^ in BT20 cells to p = 1.53×10^-209^ in SAOS2 cells, by Wilcoxon non-parametric test (**Figure S2 A-V**). IOMM (MNG) was the only exception (p = 0.24) (**Figure S2 W**), likely due to reliance on a relatively small (n=106 samples) cohort collected at Columbia—due to lack of high-quality meningioma cohorts in the literature (**Table S2**)—which barely meets the minimal requirements for ARACNe analysis.

Consistent with the fact that several drugs may be virtually inert in a specific cellular context—*e.g.*, due to baseline inactivation of their high-affinity target—several cell lines presented violin plots with a clear bi-modal structure, including a mode corresponding to significant bioactivity (higher average NES value) and one showing virtually no bioactivity (average NES = 0). Strong correlation of VIPER-assessed bioactivity and reproducibility was already shown^25^.

### Drug Similarity Assessment

Consistent with these results—and based on prior studies showing the relevance of protein activity analysis in predicting drug sensitivity^12,24,25^—we analyzed the drug-specific and cell-specific protein activity profiles to assess drug eMoA and polypharmacology.

Specifically, to assess whether two drugs, *Rx_A_* and *Rx_B_*, have similar eMoA in a specific cell line *C*, we introduce a Drug Similarity Score (*DSS_A,B_*), based on the overlap of differentially active proteins in a specific cell line following treatment with the two drugs (STAR Methods). Since the eMoA of a drug is most often tissue context specific, we also define a cell line-specific eMoA similarity score *CMSS_c1,c2_* (STAR methods)

### Pleiotropy correction

Straightforward use of the DSS and CMSS metrics may result in poor specificity, because—due to non-specific engagement of cell stress or cell death pathways, for instance—unrelated drugs may still produce partially overlapping differential protein activity signatures. While using an EC_20_ concentration largely mitigates the issue, non-specific signatures still emerged as a confounding factor, leading to critical eMoA specificity loss. To mitigate this effect, we assigned a cell line-specific weight *w_i,j_* E [0,1] to each protein in the analysis—with i and *j* identifying the protein and cell line, respectively—representing the overall conservation of the protein’s differential activity across all perturbational profiles for that cell line. The weight is computed using a sigmoidal transformation of the absolute NES value, for a specific protein and cell line, across all drugs treatments. Thus, proteins similarly affected by many drugs will produce lower (weighted) contribution to eMoA similarity (STAR methods).

### Cellular Context Specificity

Previous perturbational profile-based studies either assessed compound activity in a single cell line or integrated it across all cell lines perturbed with that specific compound^42^. As previously suggested, however, ignoring the tissue-specific nature of a compound’s eMoA can also confound the analysis^43^. Indeed, PanACEA profiles show that individual drugs may present highly similar eMoA in a cell line subset and yet virtually orthogonal eMoA in others. For instance, as expected, resminostat and panobinostat—a class I/IIb and pan histone deacetylase (HDAC) inhibitor, respectively—present highly significant eMoA similarity in both BT20 and HCC1143 TNBC cells (p = 1.6×10^-9^ and p = 1.1×10^-12^, respectively) (**Figure 2A & Figure 2B**). In contrast, pictilisib (class I PI3K inhibitor) and neratinib (ErbB family inhibitor) have highly conserved eMoA in BT20 (p = 9.6×10^-15^) but not in HCC1143 cells (p = 0.53) (**Figure 2C & Figure 2D**), even though both represent triple basal-like breast cancer.

**Figure 2:**
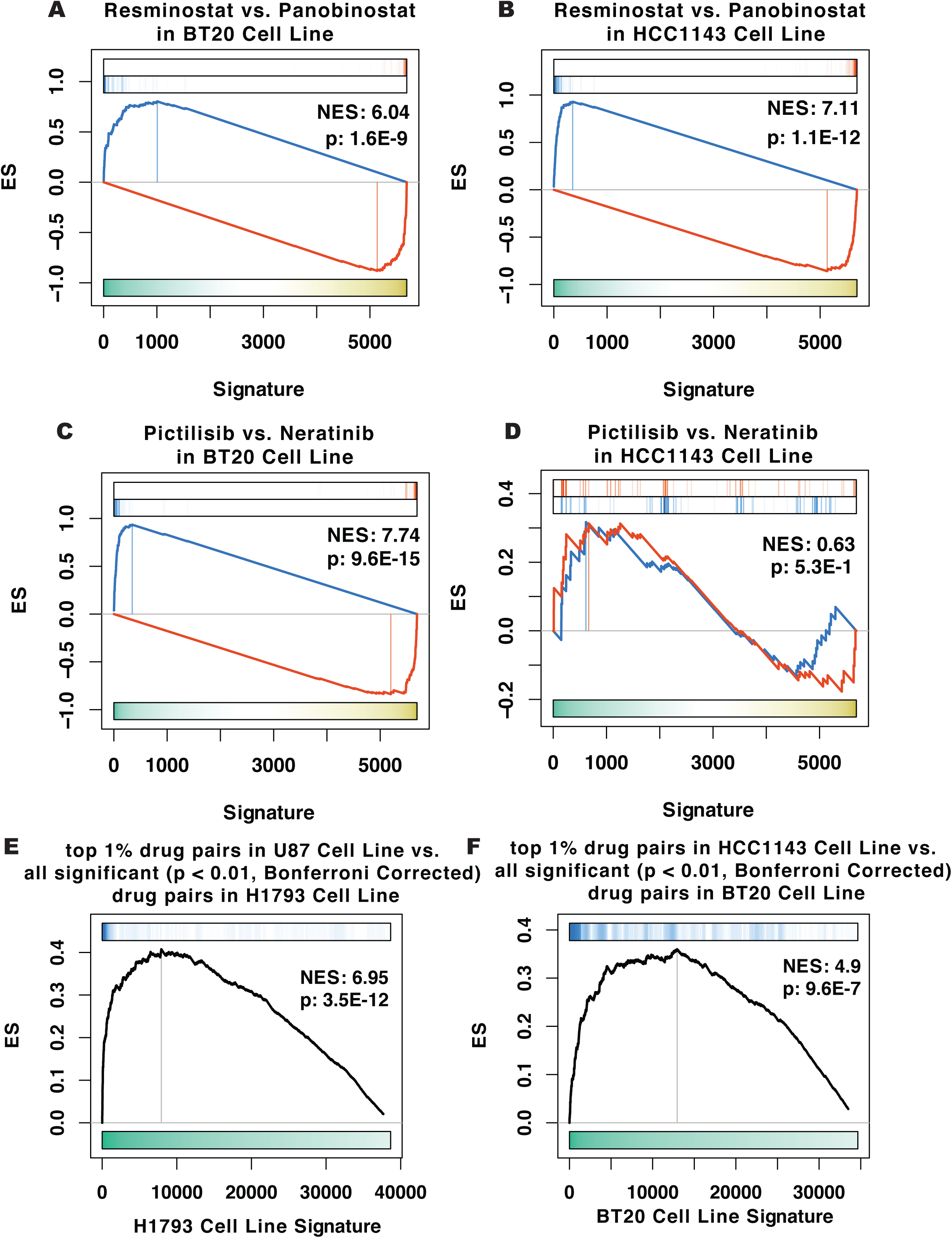
Enrichment analysis shows drug effects are cellular context dependent. (A) Two-tailed GSEA analysis shows high eMoA similarity for resminostat and panobinostat in BT20 cells. Red and blue bars represent the 25 proteins most activated and 25 most inactivated in resminostat vs. vehicle control-treated cells. The x-axis shows proteins ranked by their differential activity in panobinostat vs. vehicle control-treated BT20 cells. (B) GSEA analysis shows high eMoA conservation for resminostat and panobinostat in HCC1143 cells. (C) GSEA analysis shows unexpectedly high eMoA conservation for pictilisib (Gdc0941) and neratinib—two drugs with very different high-affinity binding targets—in BT20 cells. (D) GSEA analysis shows low eMoA conservation for pictilisib (Gdc0941) and neratinib in HCC1143 cells. (E) GSEA analysis of the 1% of drug pairs with the highest eMoA similarity in U87 cells vs. all statistically significant (p < 0.01, Bonferroni corrected) drug pairs in H1793 cells, ranked by their p-value, shows high drug eMoA conservation, despite the different cell line histology (glioblastoma vs. lung adenocarcinoma, respectively). (F) The same analysis for HCC1143 vs. BT20 cells (Both triple negative breast cancer) shows 5 orders of magnitude lower eMoA similarity compared to U87 vs. H1793.

To further explore the tissue-specific nature of drug activity, we assessed the enrichment of drug-pairs with the highest eMoA conservation in one cell line (top 1% DSS score) in drug-pairs ranked by their DSS score in a second cell line. Perhaps surprisingly, comparison of histologically distinct cell lines, such as U87 (GBM) and H1793 (LUAD), showed strikingly significant eMoA conservation across a wide range of drugs (p = 3.5×10^-12^) (**Figure 2E**). In sharp contrast, histologically related cell lines, such as HCC1143 and BT20, both TNBC-derived, presented 5 orders of magnitude lower enrichment statistics (p = 9.6×10^-7^) (**Figure 2F**).

These results suggest that eMoA analysis must directly account for tissue-specificity, independent of cell histology. Indeed, averaging results across multiple cell lines (including histology-matched ones) only diluted eMoA similarity assessment, thus failing to reveal highly conserved yet subset-specific mechanisms. This is reasonable, since different cell lines may harbor different mutations—potentially bypassing or abrogating drug activity—or represent distinct molecular states—where primary or secondary targets of the drug may or may not be expressed. A comprehensive map of such cell specific drug eMoA differences is still missing. For instance, the estrogen receptor (ESR1), an important pharmacological target in luminal breast cancer, has low or no expression in several PanACEA cell lines. As such, conservation of ESR1 inhibitors’ eMoA in these cells is likely to be mediated by off-target effects. Yet since drug targets—both high and low-affinity—are only partially characterized, the cell lines in which a drug eMoA is conserved cannot be selected *a priori*.

Another advantage of using multiple cell lines is to mitigate the potentially high false negative rate of computational methodologies designed to effectively minimize false positives. As such, eMoA conservation in a subset of cell lines may help identify mechanisms that are far more broadly conserved, as also confirmed by our validation assays.

To address eMoA context specificity in unbiased, data-driven fashion, we performed cluster analysis of each drug’s eMoA as assessed in each of the 23 available cell lines, thus revealing cell line subsets (clusters) presenting virtually identical intra-cluster yet orthogonal inter-cluster eMoA (STAR methods). For instance, consider drugs such as crizotinib, imatinib, nilotinib, tamoxifen, topotecan and vandetanib, all reported as having rather distinct high-affinity targets (**Figure S3; Table S3).** Cluster analysis shows that the eMoA of these drugs is highly conserved within each of two to three cell line clusters (positive DSS) yet virtually inverted across clusters (negative DSS). Notably, emergence of only two to three conserved-eMoA clusters, on average, across 23 cell lines suggests that only a few distinct, context-specific mechanisms exist for each drug, thus making the problem tractable.

### Mechanisms affecting drug eMoA

To address whether drug eMoA may be associated with mechanistic clues—thus providing potential biomarkers of drug response—we asked whether specific mutations, as reported in DepMap^44^, were over-represented within individual eMoA clusters, as detected by the analysis (see STAR Methods). A potential issue is the partial overlap (n=14) between PanACEA and DepMap cell lines, which reduces the statistical power of the analysis. As a result, statistically significant mutational enrichment (p < 0.05, FDR corrected) only emerged for 10 of the tested drugs (**Table S4**), suggesting that, as PanACEA increases in size, additional mechanisms may be revealed.

Among these, mutations in TRRAP (a histone acetyltransferase adaptor) were significantly enriched in cluster 4 of the mitochondrial metabolism drug Cpi613, an established tricarboxylic acid (TCA) cycle enzymes inhibitor, including the TRRAP substrate Acetyl-CoA. Similarly, eMoA cluster 3 of entinostat—a class I selective HDAC inhibitor— was enriched in mutations affecting chromatin remodeling genes, including the histone acetyltransferase (HAT) CREBBP (a COSMIC^45^ Tier 1 oncogene) and TRRAP. This is consistent with the key role played by histone acetylation homeostasis in modulating this drug’s eMoA. Statistically significant mutations for this drug were identified in additional COSMIC Tier 1 genes, such as KMT2C and RAD50.

To increase statistical power, we assessed mutation enrichment across multiple mechanism-related drugs (see STAR Methods). As an illustrative example, consider the role of TP53—the most frequently mutated tumor suppressor in human cancer—on the relatively large number of chemotoxic drugs in PanACEA. As expected, the analysis confirmed significantly greater similarity in drug eMoA for TP53^Mut^ vs. TP53^WT^ cell lines, suggesting that somatic TP53 mutations effectively “re-wire” the eMoA of chemotoxic agents. Similar results emerged for other drug classes, including HDAC, PI3K and EGFR inhibitors (**Figure S4**). Extension of these analyses to all drug classes and mutations in DepMap is summarized in **Table S4**.

### Drug eMoA Similarity Network (eMoA Network)

Even though a subset of high-affinity binding targets of PanACEA drugs may be found in existing databases (*e.g.*, DrugBank^46^), their proteome-wide activity and polypharmacology effects are still largely uncharacterized. In particular, DrugBank does not include potential high- and medium-affinity binding targets that cannot be revealed by traditional *in vitro* assays and completely ignores secondary, tissue specific effectors.

To systematically elucidate the tissue-specific mechanisms of tested drugs, we leveraged the previous section’s analysis to identify eMoA conservation at the level of each individual cell line cluster. Specifically, we leveraged hierarchical clustering to identify cell line subsets (clusters) with statistically significant intra-cluster drug eMoA conservation, as assessed by the CMSS metric. Since clusters with statistically significant conservation for two different drugs (*Rx_A_* and *Rx_B_*) may have variable overlap, we only considered cluster pairs with ≥ 50% overlap (*i.e.*, Jaccard index ≥ 1/3) and with ≥ 2 cell lines each (STAR methods).

Drug pairs (*Rx_A_* and *Rx_B_* ) in each conserved eMoA cluster are then connected by an edge, resulting in a global eMoA similarity network (henceforth, eMoA network). Each edge is assigned a p-value by integrating the DSS p-value for each cell line in the conserved cluster using Stouffer’s method (see STAR methods for a relevant example). Visualizing this network is challenging due to the large number of edges; we thus report it in tabular format (**Table S5;** STAR methods). For visualization purposes, however, we generated a simplified eMoA network with an edge connecting all drug pairs with statistically significant eMoA conservation in one or more cell line clusters and an associated p-value reflecting the most significant cluster-specific edge for that pair.

### eMoA Network Modularity Analysis

Network modularity analysis—such as used to identify protein complexes in protein-protein interaction network^47–52^, for instance—represents an effective mean to identify subset of drugs (modules) presenting high intra-module and low-extra-module eMoA conservation. In addition, such analyses can be used to identify drugs presenting strong activity conservation across two or more modules, thus revealing potential polypharmacology. For this purpose, we leveraged the ClusterONE algorithm^53^, followed by a conservative statistical significance analysis (STAR methods). The analysis identified 30 distinct drug modules, associated with equally distinct pharmacologic properties (**Figure 3A & Table S6**).

**Figure 3:**
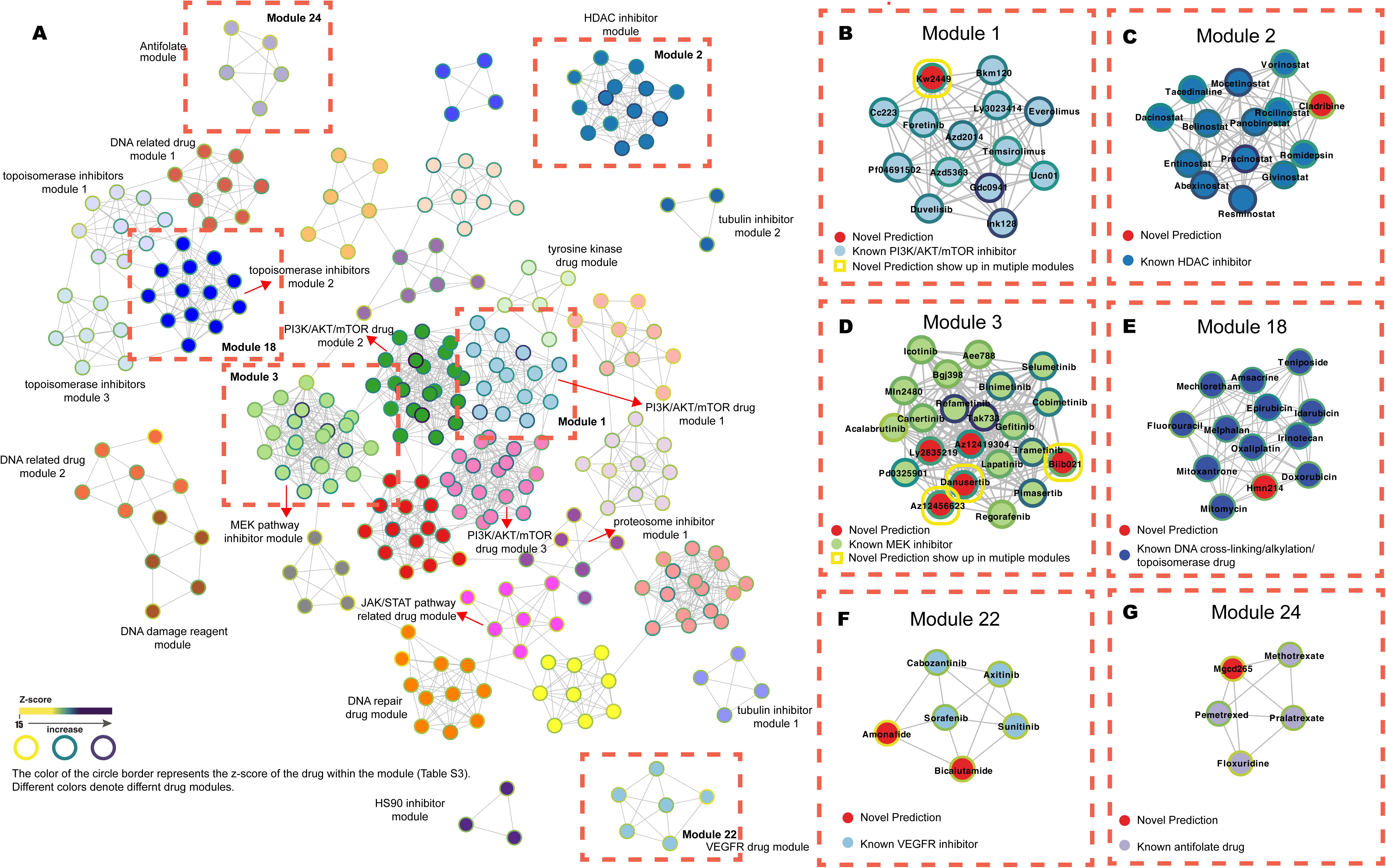
PanACEA derived drug functional map with highlighted examples. (A) In the 30 drug modules identified by eMoA modularity analysis, each node (circle) represents a drug, with same-color nodes indicating assignment to the same module and circle color representing the z-score of the Drug Specificity Test (*i.e.*, the confidence in drug-module assignment) (STAR Methods, **Table S3**). Modules with a clearly identifiable established MoA are labeled accordingly (*e.g.*, tubulin inhibitor module 2). Gray lines identify drug pairs with significant eMoA similarity in at least two of the 23 cell lines. Six highly representative modules, identified by red dotted boxes, were selected for detailed discussion and validation and are illustrated with further details in panels (B-G). Within each panel, same-color nodes indicate drugs with an established consensus MoA (*e.g.*, PIK3/AKT/mTor inhibitors). Red nodes identify drugs not previously known to share the module’s consensus MoA and thus representing either novel MoA or polypharmacology candidates. A yellow rounded-edge square identifies drugs shared by multiple modules.

To assess intra-module eMoA conservation, we computed a Drug Modularity Score (DMS), and its associated p-value, representing the overall confidence that a specific drug belongs to a module (STAR Methods).

Based on literature analysis, most drug modules were highly enriched in drugs with the same established MoA (i.e., related high-affinity targets). However, several drugs were placed in multiple modules, both related and unrelated to their established MoA, thus suggesting potential polypharmacology. This may either be a direct effect, based on a drug’s off-targets, or an indirect one, due to tissue-specific crosstalk or canalization mechanisms in a subset of the cell lines. Some key examples are shown in **Figure 3**. Comprehensive validation of all novel eMoA predictions is prohibitive, but we illustrate examples for some of the more relevant modules, including M1, M2, M3, M18 and M22, and then move to validate polypharmacology and/or novel eMoA predictions of selected compounds.

### Module M1

13 of 14 drugs in this module represent PI3K/AKT/mTOR axis inhibitors in Drug Bank (**Figure 3B)**. The remaining drug, Kw2499, is reported as a multi-kinase inhibitor, whose high-affinity binding targets include FLT3, ABL, ABL-T315I and Aurora kinase^54^. eMoA analysis, however, shows Kw2449 as having highly significant DSS—ranging from 10^-27^ to 10^-326^—with the other 13 drugs, critically across a majority of cell lines (13 of 23 in the study). This cannot be accounted for as an indirect effect mediated by its primary targets, because no other FLT/ABL/AURKA inhibitor in PanACEA was found in this module (*e.g.*, ENMD-2076).

### Module M2

13 of 14 drugs in this module are established HDAC inhibitors (**Figure 3C**). The remaining one, cladribine, is an established adenosine analog that prevents the adenosine deaminase enzyme from modulating DNA processes^55^. Remarkably, its DSS with belinostat and rocilinostat—both established HDAC inhibitors—is even more significant (p ≤ 1.22×10^-136^ and p ≤ 1.2×10^-105^, respectively), than the DSS between Belinostat and Rocilinostat (p ≤ 5.21×10^-99^), as supported by three distinct cell lines (PANC1, HSTS and GIST430). While the eMoA of cladribine may suggest potential functional similarity to HDAC inhibitor-mediated chromatin remodeling, other adenosine and, more generally, purine analogs in PanACEA—*e.g.*, pentostatin, clofarabine, mercaptopurine, and thioguanine—presented no significant DSS with HDAC inhibitors, suggesting a potentially unique moonlighting function for this drug.

### Module M3

17 of 22 drugs in this module represent MAPK pathway inhibitors, including inhibitors of MEK and of its upstream modulators and downstream effector kinases (**Figure 3D**). The remaining 5 drugs represent a quite heterogeneous mixture, including Az12419304 and Az12456623—two small molecule inhibitors of unknown function—biib021 (an HSP90 inhibitor^29^), danusertib (an AURKA inhibitor^56^), and ly2835219 (a CDK4/6 inhibitor^57^).

### Module M18

12 of 13 drugs in this module represent DNA cross-linking/alkylation and topoisomerase inhibitors (**Figure 3E**). The polo kinase (PLK1) inhibitor hmn214^58^ was also predicted as a highly statistically significant member of M3 (ranging from p ≤ 5.26×10^-42^ with irinotecan to 1.49×10^-107^ with fluorouracil), potentially consistent with its role as a modulator of DNA damage repair mechanisms.

### Module M22 (Figure 3F)

4 of 6 drugs in this module (sunitinib, axitinib, sorafenib, and cabozantinib) represent VEGF pathway inhibitors. The remaining two drugs include amonafide—a topoisomerase inhibitor^59^ that, surprisingly, failed to cluster with other topoisomerase inhibitors in M22—and bicalutamide, a selective androgen receptor blocker^60^. Both had highly significant DSS with VEGF pathway inhibitors. Specifically, amonafide’s eMoA similarity with sorafenib and cabozantinib was highly significant (p ≤ 1.03×10^-34^ and p ≤ 6.22×10^-55^, respectively), while bicalutamide similarity with axitinib and sorafenib was even more significant (p ≤ 1.36×10^-72^ and p ≤ 4.64×10^-92^, respectively).

### Module M24

3 of 5 drugs in this module (methotrexate, pemetrexed, and pralatrexate) represent established antifolate compounds (**Figure 3G**). A fourth one (floxuridine) is an established inhibitor of thymidylate synthetase^61^, an enzyme playing a key role in folate biosynthesis. The final drug (Mgcd265, also known as glesatinib) is an established multi-target kinase inhibitor^62^, with no known effect on folate biosynthesis. This is especially relevant as there are no other examples of drugs with polypharmacology activity in this pathway.

Critically, identification of drug-class specific modules could not be achieved by correlating drug sensitivity profiles from large scale resources such as DepMap (**Figure S5**), suggesting that a more mechanism-based approach, such as the one implemented via protein activity analysis, is required for effective eMoA similarity assessment.

### Tissue Specificity

Modularity analysis also highlighted the tissue specific nature of compound eMoA. For example, Module **M29** includes four DNA-damaging drugs (mercaptopurine, ifosfamide, streptozotocin, and procarbazine) and two steroidogenesis inhibitors (mitotane and aminoglutethimide) (**Figure S6A)**. While two of the cytotoxic drugs ifosfamide and mercaptopurine showed highly similar activity across most cell lines, aminoglutethimide, mitotane, ifosfamide, procarbazine and streptozotocin formed a highly statistically significant sub-module only in three sarcoma-derived cell lines, including SAOS2 and MSKRMS (osteosarcoma) and SKNEP (Ewing sarcoma), again providing a potential sarcoma-specific link between steroidogenesis and DNA-mediated cytotoxicity. An additional compelling example is provided by Module **M13** (**Figure S6B)**, where drugs have highly distinct established MoA. While individual, statistically significant similarity, represented by network edges, are only supported by a handful of tissues, potentially also due to false negatives, a highly interconnected sub-module, comprising five drugs (gmx1778, bay-117085, cp100356, ym201636 and paroxetine) emerged as highly conserved in the two TNBC cell lines (HCC1143 and BT20) suggesting a mechanism of drug effect canalization that is specific to this tumor subtype. These illustrative examples show that tissue-specificity plays a critical role in determining the overall effect of drugs on the proteome, suggesting interesting tissue-specific effects for future investigation.

### Experimental and Structure-based Validation of Drug Mechanism of Actions and Polypharmacology

To mitigate the time and labor-intensive nature of experimental eMoA validation, we focused on three modules comprising extensively characterized compounds. These were used to validate different aspects of our predictions including (i) within-class polypharmacology (***i.e.***, compounds predicted to target the same protein class, *e.g.*, protein kinases), (ii) cross-class polypharmacology (i.e., compounds predicted to target different protein classes), and (iii) *de novo* elucidation of uncharacterized compound mechanism. For these purposes, we measured transcript and protein abundance of critical activity reporters to detect functional effects and coupled them with proteome-wide mass spectrometry-based PISA assays and structure-based analyses to further elucidate their mechanistic rationale. Notably, while compound proteome-wide eMoA was predicted based on perturbation at the EC_20_ concentration, functional and mechanistic assays were performed at higher concentration to ensure a more detectable biochemical effect (see STAR Methods).

### In-class polypharmacology

Kw2449—an ABL/FLT3 and Aurora kinases inhibitor—co-clustered with established mTOR pathway inhibitors in multiple cell lines (module **M1**), including SAOS2. We thus proceeded to measure the ratio of phosphorylated (p-P70S6K) p70 (ribosomal S6 kinase) vs. GAPDH in drug vs. DMSO in SAOS2 cells, by Western blot (WB). Similar to treatment with temsirolimus—an established mTOR inhibitor as positive control—p-70S6K levels were significantly reduced in kw2449 treated cells compared to vehicle-treated and negative control cells (mTOR pathway activation with 20% FBS), (**Figure 4A-C**). To elucidate potential mechanisms, we performed PISA assays to detect biochemically engaged c proteins^30^. The assay successfully identified kw2449’s canonical targets, including AAK1, AURKA, and TBK1, among others (**Figure 4D**). However, they also revealed RPS6KB1 (P70S6K) as an additional stabilized protein (log_2_ fold change ***FC*** = 0.27, p = 1.09 × 10^-5^, *p-values are FDR corrected, unless explicitly stated).* This kinase plays a key role in mTOR pathway regulation, thus providing a mechanistic rationale for the observed functional effects. Docking simulations^33,63^ starting from poses derived from the PrePCI database^31^ suggest a Kw2449/P70S6K interaction is likely mediated by the insertion of the compound’s purine ring into the active site of P70S6K (**Figure 4E**), with an associated binding free energy of -9.8 kcal/mol, as assessed by Induced Fit Docking (IFD) (STAR Methods) (**Table S7 H**). PrePCI (STAR methods), provided additional clues. Specifically, this structure-based algorithm recapitulated established Kw2449 targets but also provided high-confidence predictions (FPR < 0.001) for several additional proteins in the PI3K/AKT/mTOR pathway^64^ (**Table S7 A)**, including INPP5D (also known as SHIP1). INPP5D is the phosphatidylinositol 3,4,5-trisphosphate 5-phosphatase-1 responsible for hydrolyzing the 5-phosphate of the PI(3,4,5)P3 lipid to produce PI(3,4)P2. Both lipids are potent mitogenic signals involved in AKT activation^65^. Taken together these data confirm the predicted role of Kw2449 in modulating PI3K/AKT/mTor pathway signals, as potentially mediated both by direct interaction with the P70S6K kinase as well as by the potential interaction with additional members of this pathway, including INPP5D, whose off-target inhibition would also contribute to p-70S6K levels reduction.

**Figure 4:**
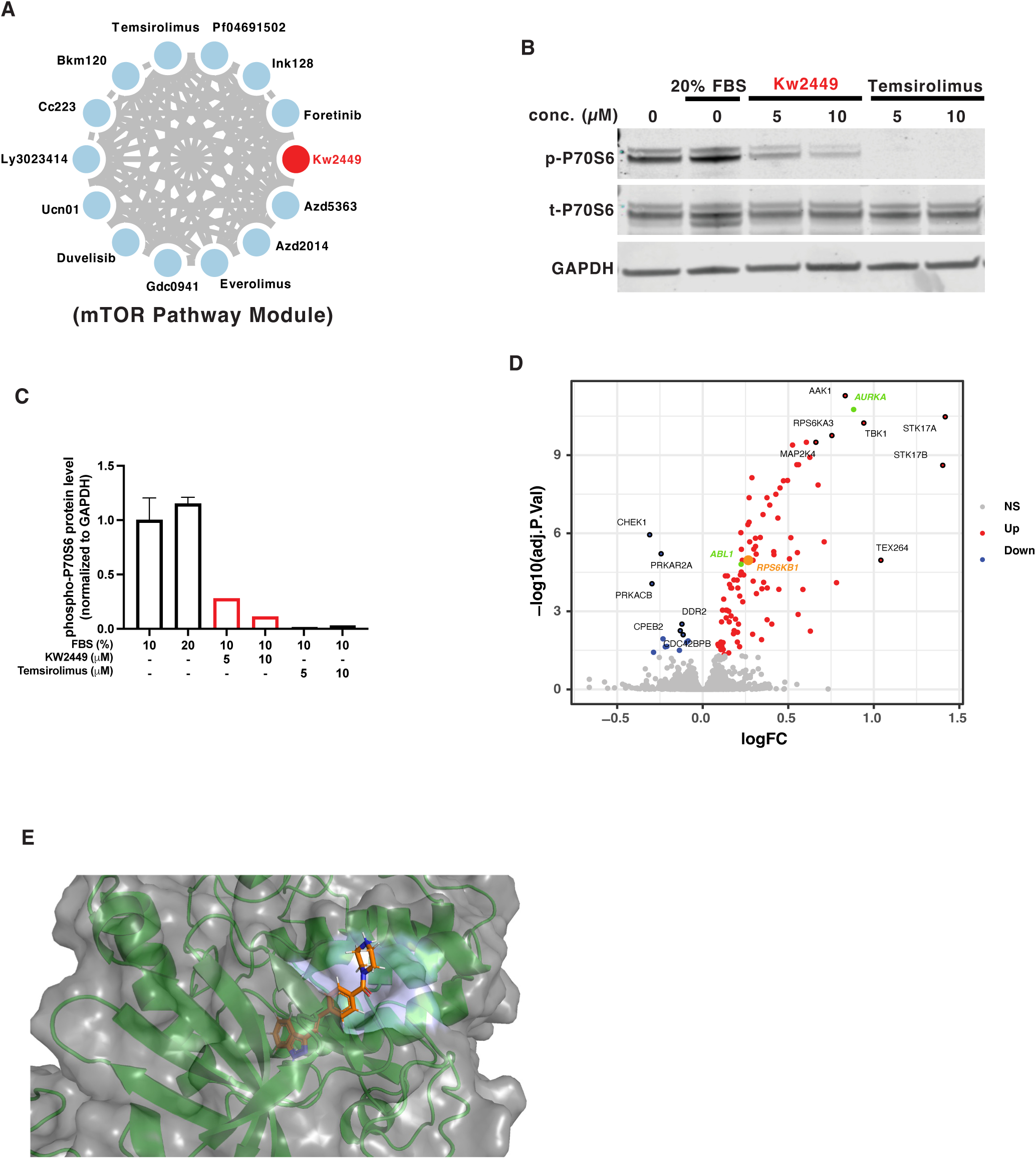
Validation of polypharmacology prediction for mTOR module. (A) In the PI3K/AKT/mTOR pathway inhibitor consensus module (M1), red circles identify the drugs selected for experimental validation. (B) Consistent with predictions, Kw2449 inhibited phosphorylation of the canonical mTor substrate p70 ribosomal S6 kinase (P70S6K) in dose-dependent manner. Saos2 cells were cultured in 10% FBS, to induce mTor pathway activation, and then treated for 6 hours with vehicle control or the indicated concentration of Kw2449 and temsirolimus (a known mTor inhibitor). Cell pellets were collected and lysed and even amounts of proteins were separated by SDS-PAGE and blotted for phospho-P70S6K (p-P70S6), total-P70S6K (t-P70S6) and GAPDH as loading control. (C) Quantification of treatment-mediated p-P70S6K abundance changes were normalized to GAPDH. (D) In the PISA assay results for Kw2449, green colored proteins are its known binding targets, orange-colored proteins are novel target proteins. PISA detected that Kw2449 stabilizes RPS6KB1 (protein name: P70S6K), which suggests a target engagement between Kw2449 and P70S6K. (E) Atomic-level representation shows the protein-compound-interacting region, based on a Kw2449-P70S6K structural model.

### Cross-class polypharmacology

Mgcd265—a MET/AXL kinase inhibitor—was predicted as a folate metabolism inhibitor in multiple cell lines including LoVo (**Module 24)**. Treating these cells for 24h, at various compound concentrations (0.5 μM, 1 μM and 5 μM), decreased DHFR protein levels at all three concentrations (**Figure 5A-C**). Using thermal shift assays, we observed significant stability changes consistent with MGCD265 engagement of its known targets (including MET, ABL1, and TBK1; **Figure 5D**). However, GSEA analysis of proteins exhibiting significant stability changes in the thermal shift assay (ranked by p values, p < 0.01, FDR corrected) also identified the GPI transamidase (GPIT) CORUM complex^66^ as the top hit (p = 1.01 × 10^-4^, **Figure 5E**). Moreover, the “GPI anchor pathway” was also ranked as one of the top pathways in KEGG^67^, Reactome^68^ and WikiPathways^69^ by GSEA analysis (**Figure S7**), with four GPIT complex subunits significantly destabilized by mgcd265 (**Figure 5D**), including PIGT (FC = -0.72, p = 2.97 × 10^-9^), PIGS (FC = -0.39, p = 2.01 × 10^-7^), PIGU (FC = -0.5, p = 1.1 × 10-^-3^) and PIGK (FC = -0.46, p = 1.96 × 10^-3^). The GPIT complex plays a key role in anchoring the folate receptor (FR) to the cell’s membrane to support folate uptake^34^.

**Figure 5:**
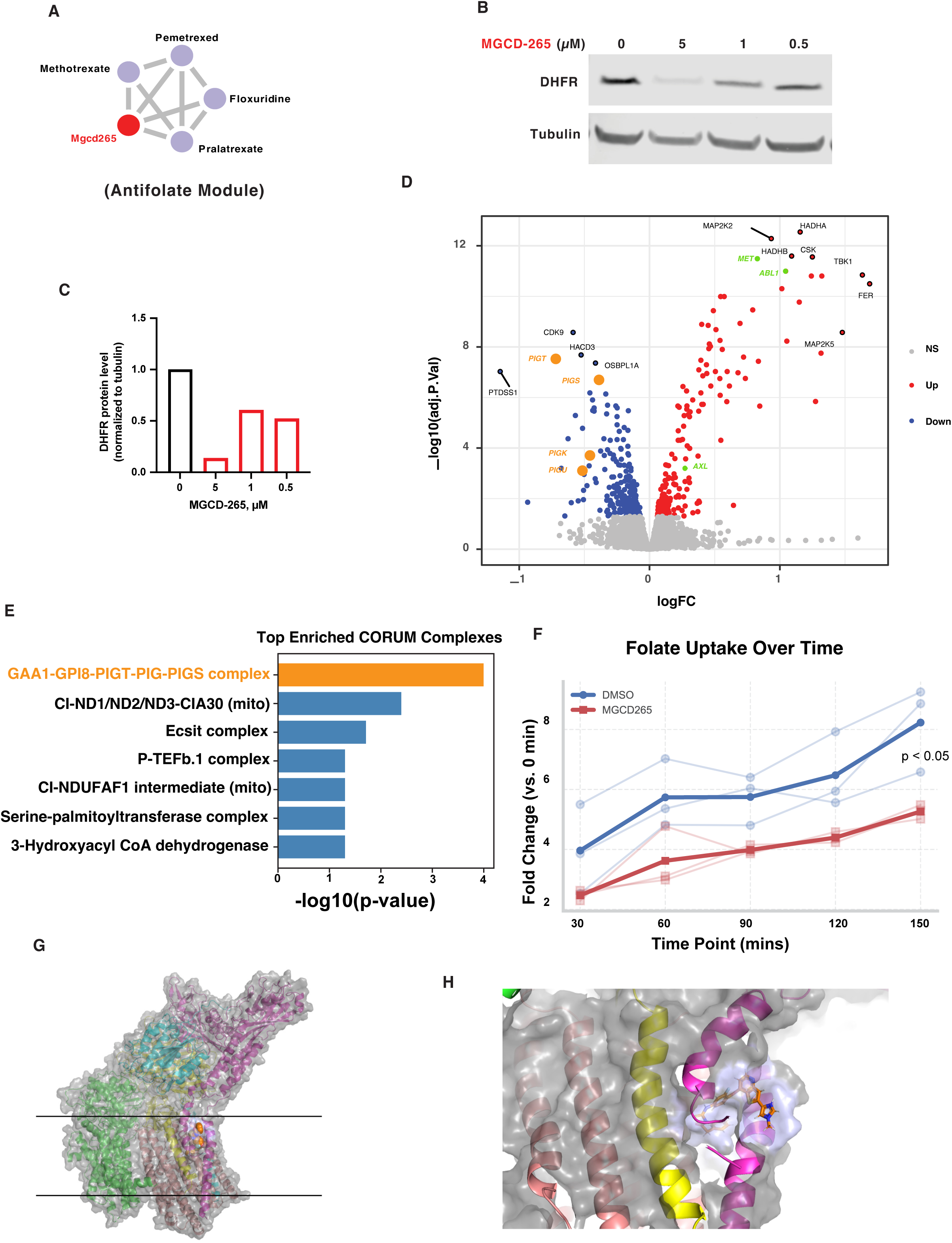
Validation of novel drug polypharmacology prediction for folate metabolism module. (A) In the antifolate consensus module (M24), red circles identify the drugs selected for experimental validation. (B) Consistent with predictions, Mgcd265 induced DHFR protein abundance reduction. LoVo cells were treated with Mgcd265 for 24 hours, at the indicated concentrations. Cell pellets were collected and lysed, and even protein amounts were separated by SDS-PAGE and blotted for DHFR and tubulin (loading control). (C) Quantification of treatment-mediated DHFR abundance changes was normalized to tubulin. (D) In the PISA assay result for Mgcd265, green colored proteins are its known binding targets, and orange-colored proteins are novel target proteins. (E) Gene set enrichment analysis for identified proteins (p < 0.01, FDR corrected) from Mgcd265 PISA assays against CORUM complexes database. (F) Folate uptake assay was performed for Mgcd265 treated cells vs. DMSO treated cells. (G) Atomic level representation shows the GPIT structural model with bound Mgcd265 in the endoplasmic reticulum (ER) membrane region depicted within the black lines. (H) The protein-compound-interacting region of Mgcd265 is highlighted.

Structure-based Induced Fit Docking (IFD) simulations^33^ indicates that the interaction between mgcd265 and the GPIT complex is predicted to occur in the transmembrane region of the GPIT complex (**Figure 5G & H**), with a predicted binding free energy of -15.6 kcal/mol (**Table S7 H**). We thus hypothesized that mgcd265 may destabilize the GPIT complex and prevent effective FR anchoring, thus reducing folate uptake. Consistent with this, EC-17 uptake assays showed statistically significant reduction in folate uptake in drug vs. DMSO treated cells (p < 0.05, **Figure 5F**). Since thermal shift assays may reflect either direct engagement or downstream effects on complex assembly, these results do not necessarily prove direct binding. However, taken together, the repression of DHFR protein abundance, disruption of the GPIT folate anchoring complex, and IFD-based analyses strongly support the prediction of mgcd265 as novel folate metabolism inhibitor.

### De novo elucidation of compound pharmacology

finally, the proposed framework can help characterized compounds with unknown MoA. Module **M3** is almost exclusively comprised of established MEK inhibitors, except for biib021 (an HSP90 inhibitor) and Az12419304 (unknown MoA). To validate these predictions, we measured the ratio of phosphorylated (p-Erk) to total Erk (t-Erk) in drug vs. DMSO-treated N87 cells by WB. This ratio was significantly reduced by both drugs, in dose-dependent manner (**Figure 6A-C**). Focusing specifically on Az12419304, PISA assays showed that BRAF—a direct upstream MEK regulator—was stabilized by Az12419304 (FC = 0.88, p = 3.21×10^-8^, **Figure 6D**), consistent with its predicted role as MEK pathway inhibitor (**Figure 6B**). PrePCI analysis (**Table S7 C**) confirmed BRAF as a top hit for Az12419304. In addition, MAPK14 (FC= 2.18, p = 2.68×10^-11^) and MAPK11 (FC = 1.60, adjusted p = 2.86×10^-6^) also emerged as top stabilized proteins by Az12419304 in PISA assays. Finally, EPHA2 also emerged as a top hit by PISA (FC= 0.75, p = 1.2×10^-9^) and was among the top cognate binding partner of the compound as predicted by PrePCI (**Table S7 C**). Finally, induced fit docking simulations suggest that the Az12419304/BRAF interaction is likely mediated by insertion of the compound into the active site of BRAF (**Figure 6E**), with a predicted binding free energy of -12.7 kcal/mol (**Table S7 H**). PrePCI also identified several high-confidence predictions for **M3** drugs as MAPK pathway (GO: 0000165) inhibitors (**Table S7 C-F**), including predictions supported by direct experimental evidence in PDB, such as danusertib/EPHB2 and ly2835219/CDK6. Further analysis revealed that its statistically significant effectors, as assessed by PISA assays (p ≤ 0.05, FDR corrected), were highly enriched (p = 2.4×10^-5^) in proteins predicted to have the highest interaction likelihood by PrePCI analysis (**Figure S8**); leading edge proteins from this analysis were significantly enriched in Reactome MAPK pathway members (p = 2.0×10^-3^), RHO GTPases (p = 1.0×10^-3^), receptor tyrosine kinases (p = 3.9×10^-4^), and their downstream effector AP-1 family transcription factors (p = 6.2×10^-5^). This supports the proposed concept of a field effect, whereby drug function may be mediated by the combined effect of a large number of proteins rather than just by their high-affinity target(s). See similar enrichment was revealed for biib021 (**Figure S9**).

**Figure 6:**
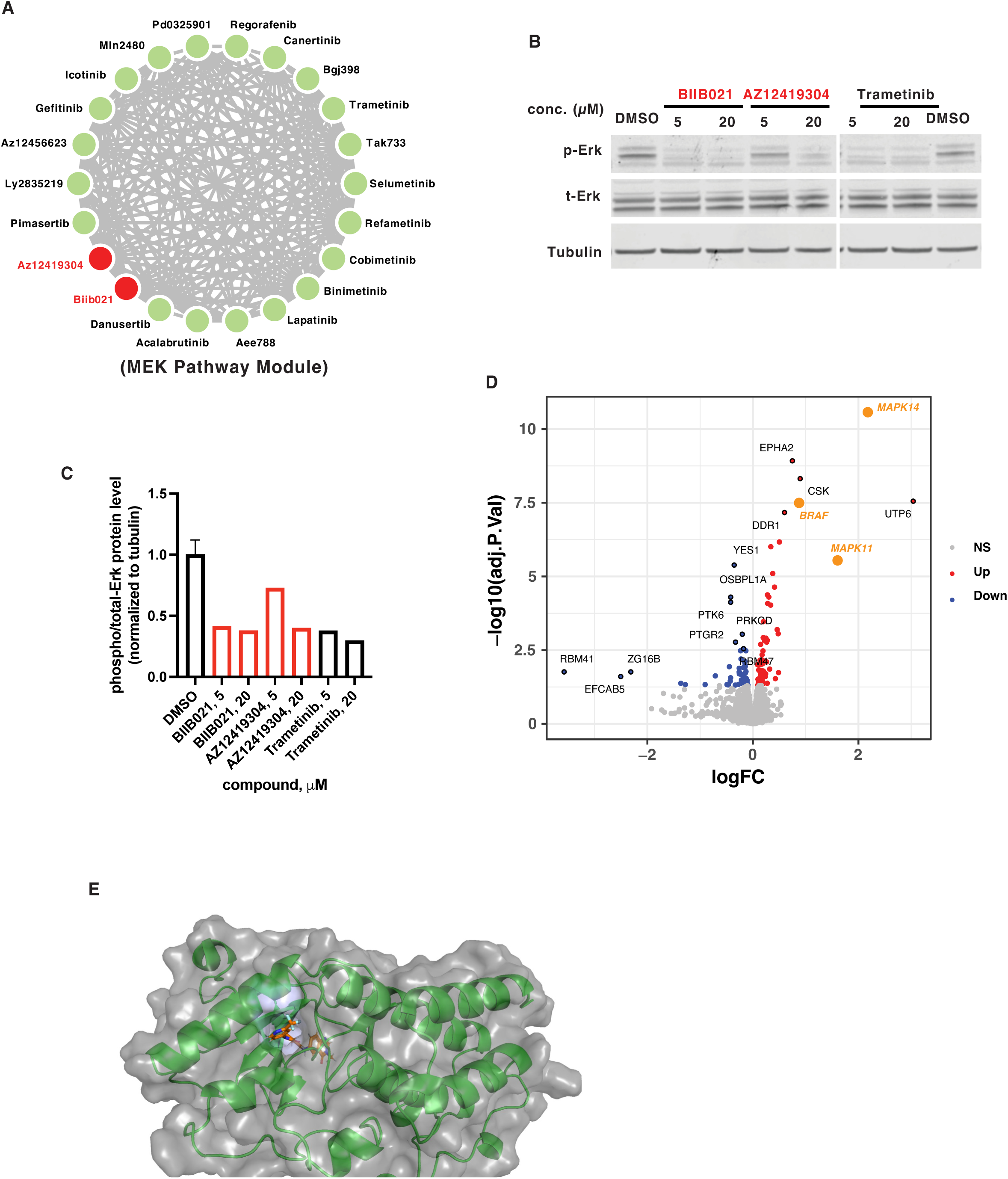
Validation of novel drug eMoA or polypharmacology prediction for MEK module. (A) In the MEK pathway inhibitor consensus module (M3), red circles identify the drugs selected for experimental validation. (B) Consistent with predictions, biib021 and Az12419304 inhibited phosphorylation of ERK, selected as the most downstream reporter of MAPK pathway activation. N87 cells were treated with biib021 or Az12419304 for 4 hours, at the indicated concentrations. Cell pellets were collected and lysed, and even protein amounts were separated by SDS-PAGE and blotted for phospho-ERK (p-ERK), total-ERK (t-ERK) and tubulin as loading control. Trametinib (a known MEK inhibitor) was used as positive control. (C) Quantification of treatment-mediated p-ERK abundance changes was normalized to tubulin. (D) The PISA assay identified Az12419304 as stabilizing BRAF, MAPK14 and MAPK11, which suggests a target engagement between them and Az12419304. (E) Atomic-level representation shows the protein-compound-interacting region based on a Az12419304-BRAF structural model.

Structure-based analyses also supported additional polypharmacology predictions that were not included in the experimental validation assays. For instance, **Figure S10** depicts a structure-based model for the complex between cladribine and HSP90AA1—a heat-shock protein included in HDAC-related pathways^64^—based on a high-confidence PrePCI prediction (FPR < 0.001). HDAC inhibitor-mediated deregulation of chaperone function has been shown to disrupt protein homeostasis and induce protein misfolding^70^ suggesting a functional link between HDAC and HSP90 inhibition. **Table S7B** shows that cladribine has PrePCI-predicted targets (FPR < 0.001) in common with five of the drugs in this module (entinostat, romidepsin, dacinostat, pracinostat, and abexinostat). In particular, all six compounds are predicted to bind HSP90AA1.

### Targeting the Undruggable Proteome

TFs have emerged as critical tumor-essential proteins in many studies. Unfortunately, most of them are also deemed undruggable for lack of specific binding pockets^71^ and structural models, usually restricted to their DNA-binding domain. We thus explored whether our framework may offer novel insight into compounds that inhibit TF activity in specific tissue contexts, either by targeting their upstream modulators or by implementing complex field-effects, due to polypharmacology.

Systematic analysis of ∼10,000 samples in The Cancer Genome Atlas (TCGA) elucidated Master Regulator (MR) proteins representing ultra-conserved, mechanistic determinants of tumor cell state, as well as key members of regulatory bottlenecks responsible for canalizing the effect of patient-specific mutations^24^. These proteins are highly enriched in essential and/or synthetic lethal proteins, thus constituting a novel class of potential drug targets^11,24,36,72,73^. Of these, 37 were recurrent (*i.e.*, identified in ≥ 15 cancer subtypes). We thus looked for drugs predicted to inhibit their activity in cell lines where they had above-average VIPER activity, compared to the average of CCLE and Genentech cell lines repositories^74^. For each MR and cell line, we show a heatmap representing the activity inhibition by the 15 top predicted drugs, as assessed by VIPER analysis of drug vs. vehicle control-treated cells (**Figure S11)**.

As expected, the three most significant inhibitors for each MR and cell line (**Figure S12**) emerged as highly cell line/tissue specific, since compound eMoA is mediated by the distinct signaling and regulatory logic of each cell line. For instance, statistically significant OVOL1 inhibitors were identified in only four cell lines. However, due to the stringency of PanACEA-based predictions, we expected individual findings to generalize to additional tissue contexts. Taken together, these data suggest that several effective, yet context-specific inhibitors of the undruggable proteome—beyond the 37 proteins studied here—could emerge.

To identify highly conserved MR inhibitors, further increasing confidence in their predictions, we show the number of cell lines where the top 15 inhibitors of each MR were identified as significant (**Figure S13**) and also summarize the analysis in a table (**Table S8-30)**.

For validation purposes, we focused on four MRs—MYC, CTNNB1, CENPF and UHRF1—as supported by key reagent availability, including optimized MR-specific antibodies^24^, cell line culture conditions, and baseline MR expression. WB assays in triplicate were used to report on the differential abundance of the target protein in drug vs. DMSO-treated controls, including within the nuclear fraction for specific TFs, such as CENPF, where the whole cell assays may be confounded by the high molecular weight and ER interactions. Results for each MR, based on p-value assessment by VIPER analysis, are discussed in the following.

1. **MYC**, one of the most frequently amplified or translocated proto-oncogenes across multiple of human maligancies^75^, still represents one of the most elusive drug targets. For this MR, we tested predicted inhibitors pimasertib and refametinib (MEK1/2 inhibitors) in HCC1143 (TNBC) (NES_pima_ = -6.2, p ≤ 5.65×10^-10^; NES_refa_ = -5.14, p ≤ 2.75×10^-7^) and LoVo (COAD) cells (NES_pima_ = -6.32, p ≤ 2.62×10^-10^; NES_refa_ = -8.23, p ≤ 2.22×10^-16^) (**Figure 7A)**.
2. **CTNNB1** (β-catenin) is a key WNT signaling effector, playing an important role in tumorigenesis^76^. For this protein, we tested the predicted DNA intercalator dactinomycin in KRJ1 cells (GEP-NET) (NES = -4.17, p ≤ 3.05×10^-5^) (**Figure 7B**).
3. **CENPF** is a large coiled-coil protein involved in centromere maintenance during mitosis^77^. In previous work, we identified CENPF and FOXM1 as critical synergistic dependencies of malignant castration resistant prostate cancer^78^ and confirmed their aberrant co-activation in several of the most aggressive TCGA malignancies^24^. For this MR we tested the topoisomerase II inhibitor amsacrine in HF2597 and U87 cells (GBM) (NES = -12.8, p ≤ 1.64×10^-37^ and NES = -12.31, p ≤ 8×10^-35^, respectively) (**Figure 7C top**) as well as the proteasome inhibitor bortezomib (NES = -11.4, p ≤ 2.64×10^-30^) (**Figure 7C bottom**). Due to the high molecular weight of this protein and its dependency on nuclear localization, we performed WB assays in the nuclear fraction of tested cells.
4. **UHRF1** plays an important role in the maintenance of DNA methylation in mammalian cells^79^ and represents a candidate drug target due to its essentiality in multiple human malignancies^80,81^. For this MR, we tested pracinostat (HDAC inhibitor) in HF2597 cells (GBM) (NES= -11.07, p ≤ 1.75×10^-28^) and bortezomib (proteasome inhibitor) in H1793 cells (LUAD) (NES = -11.21, p ≤ 3.64×10^-29^ and NES = -10.65, p ≤ 1.74×10^-26^, respectively) (**Figure 7D**).

**Figure 7:**
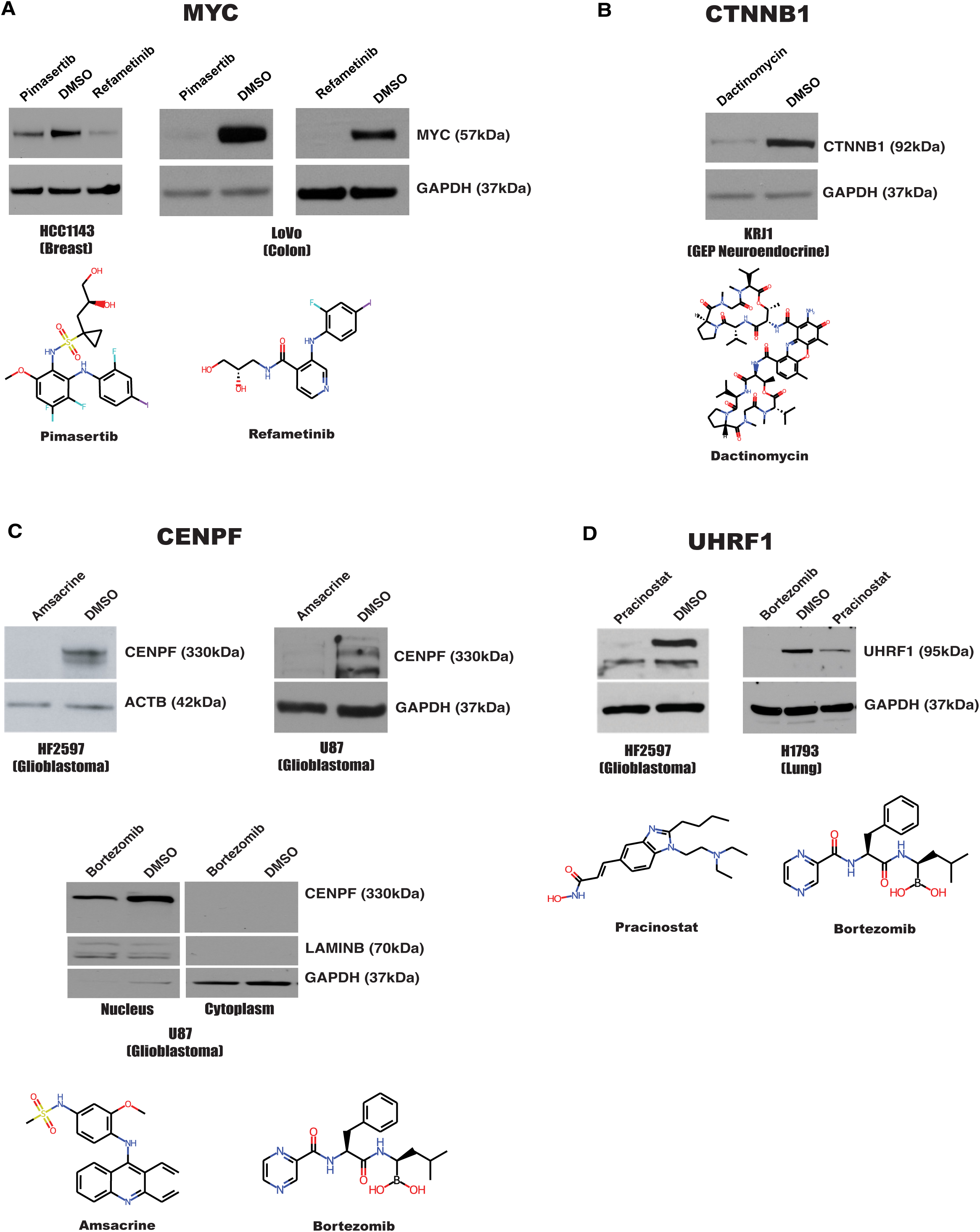
Validation of context-specific MR inhibitor predictions. (A) Western Blot (WB) assays confirm that pimasertib and refametinib reduce MYC expression in HCC1143 and LoVo cells. (B) WB assays show dactinomycin reduces CTNNB1 expression in KRJ1 cells. (C) WB assays show amsacrine reduces CENPF expression in HF2597 and U87 cells. Cellular fractionation-based WB shows that bortezomib reduction of CENPF in U87 cells is nucleus-specific. (D) WB assays show pracinostat reduces UHRF1 expression in HF2597 and H1793 cells, and that bortezomib reduces UHRF1 expression in H1793 cells.

Taken together, these assays confirmed that virtually all tested inhibitors were found to significantly abrogate protein abundance in predicted cells (low false positive rate). To also assess whether PanACEA-based predictions may generalize to contexts in which the effect was not predicted—also accounting for potential false negatives—we tested the predicted MYC inhibitor Pimasertib in IOMM (meningioma, ME) and HSTS (gastroenteropancreatic neuroendocrine tumor, GEP-NET) cells, where the effect was not predicted (NES = -0.12, p = 0.9 and NES = -0.05, p = 0.96, respectively) (**Figure S14**). These assays confirmed that predicted effects may generalize more broadly than the specific context in which they were revealed.

## Discussion

This study reports on a 10-year-long effort to create an extensive collection of gene expression and protein activity profiles for the network-based elucidation of tumor-relevant drug mechanism of action and polypharmacology (PanACEA). These profiles represent the genome/proteome-wide response of 23 cell lines—selected as high-fidelity models of molecularly distinct cancer subtypes—to clinically relevant drugs. The proposed design offers three main advantages: (a) the fully automated PLATE-Seq technology allowed systematic collection of genome-wide, highly reproducible perturbational profiles; (b) cell lines were chosen to represent clinically relevant models of human malignancies, based on the OncoMatch algorithm^10^, a critical aspect of the study since activity predictions in MR-matched cell lines were previously recapitulated *in vivo*—in drug-treated PDX models—for 15 of 18 tested drugs; and (c) drugs were carefully titrated at their highest sub-lethal concentration (EC_20_), independently assessed in each cell line, thus mitigating critical bias and confounder effects associated with activation of cell stress and death pathways, while avoiding non bioactive concentrations. Availability of genome-wide perturbational profiles, in particular, was instrumental to leveraging the VIPER algorithm to generate systematic and highly reproducible differential protein activity profiles for ∼6,500 regulatory and signaling proteins. The resulting data provides critical insight into pharmacologic drug activity—including detection of lower-affinity binding targets and cellular canalization-mediated mechanisms—thus supporting effective prediction of polypharmacology and extended Mechanism of Action (eMoA), which cannot be effectively dissected only based on large scale sensitivity assays.

Differential protein activity assessment in drug vs. vehicle control-treated cells was critical to study drug related mechanisms because most inhibitors modulate a protein enzymatic or regulatory activity rather than its abundance or the expression of its encoding gene—as shown for estrogen receptor inhibitors^6,7^, for instance. An additional advantage of VIPER analyses is their high reproducibility, which led to CLIA compliance and NY and CA Dept. of Health approval^82^ of the VIPER-based OncoTreat^25^ and OncoTarget^83^ algorithms, supporting their clinical use. Moreover, several studies have shown that VIPER-based protein activity measurements compare favorably with antibody-based approaches, including at single cell level, where gene dropout can severely compromise individual gene readouts^12,21^. Consistent with previous *in vitro*, *in vivo* and clinical studies, VIPER prediction accuracy was confirmed, since virtually all the drugs identified as potential context-specific MR inhibitors were functionally validated. However, due to potential false negatives, predicted effects may generalize more broadly than the context in which they were initially revealed, thus supporting the value of using multiple cell lines in the analysis.

Overall, the combination of the largest collection of genome/proteome-wide drug perturbation profiles and the associated analytical framework provides a valuable framework to help elucidate novel drug targets and mechanisms, which can be easily extended to study small molecule compounds and biologics in virtually any disease-related or physiological context. For instance, we recently showed that drug perturbation profiles helped identify drugs that successfully reprogrammed regulatory T cell tropism^36^. Taken together, these results suggest that VIPER-based analysis of drug perturbation profiles could help identify or develop novel, tissue-specific inhibitors for proteins representing candidate targets for tumors and other diseases. In addition, this suggests that pharmacological drug activity may be mediated by the indirect effects of multiple proteins, rather than only by their high-affinity targets.

The novel mechanism-based insights into drug polypharmacology and eMoA provided by this study critically extend the value of drug-perturbation profiles, whose VIPER-based analysis has been shown effective in predicting tumor-specific drug sensitivity^10^, including in castration resistant prostate cancer^24^, metastatic neuroendocrine tumors^25^, and diffuse midline glioma^84^. However, this does not affect the novelty of this study since prior work focused specifically on predicting tumor-specific drug sensitivity rather than proteome-wide drug mechanism of action (eMoA) and polypharmacology.

Similar to other computational approaches, PanACEA comes with some limitations, including potential false positives and false negative predictions, which represent a direct reflection of the same limitations previously reported for VIPER predictions^7^, where 20% of transcriptional regulator proteins and 30% of signal transduction proteins were shown to have poorly predictive regulons in some contexts. In addition, lineage-specific pleiotropic mechanisms—e.g., those associated with cell stress and death pathway activation only in epithelial cells—may also confound the analysis, thus failing to reveal context-specific differential protein activity. As shown, analyzing profiles from multiple cell lines mitigates this concern, supporting identification of general mechanisms by VIPER analysis in a limited context, as shown for MYC. In general, the false positive and negative rates arising from VIPER are similar to and arguably better than those reported for high-throughput experimental assays. Thus, the combination of high-throughput, VIPER-based computational prediction and low-throughput experimental validation—as harnessed in this study—is critical to the effective elucidation of relevant drug mechanisms.

Taken together, the results of this study suggest that PanACEA represents a highly generalizable methodological and experimental resource to support a more quantitative application of precision medicine principles, based on the identification of small molecules that target critical, context-specific dependencies of both human malignancies, with straightforward generalization to other human diseases. In particular, the proposed framework may help guide low-throughput experimental assays in a much more targeted and efficient way towards the validation of mechanisms that may be critical to elucidating the full spectrum of activity of a drug.

## Supporting information

star methods

## Data Availability

Drug perturbation RNA-seq data generated from this work have been deposited to the Gene Expression Omnibus (GEO: GSE275348). The data is also publicly accessible via a dedicated web portal (https://califano-lab.github.io/panacea.io/).

## Acknowledgements

We acknowledge the High-Throughput Screening and the Genome Sequencing and Analysis Facilities at the JP Sulzberger Columbia Genome Center, the Confocal and Specialized Microscopy Shared Resource (SCMSR) managed by the Herbert Irving Comprehensive Cancer Center at Columbia University and supported in part by the NIH/NCI Cancer Center Support grant P30CA13696. This work was also supported by the NCI Cancer Target Discovery and Development (CTD2) awards U01CA272610 and U01CA217858, Center for Cancer Systems Therapeutics (CaST) award U54CA274506, and the NIH Shared Instrumentation Grants S10OD032433, S10OD012351 and S10OD021764, all to A.C. The research of B.R.S. was also supported by NCI grants P30CA013696 and P01CA291697and by NIH grants S10OD020056 and S10OD030282, 5P30DK132710, and P30CA008748; B.H. was supported by NIH grant R35-GM139585.

## Open Access Statement

This content of this study is published under open access subject to licensing agreements. The authors allow researchers to distribute and use data from the original source provided it is distributed free of cost for non-commercial purposes. Any profit-making is strictly prohibited.

