## Supplementary material for "Compound Mechanism of Action and Polypharmacology can be Elucidated by Large-Scale Perturbational Profile Analysis": star methods

**Cell line selection:** Cell line transcriptional profiles for OncoMatch analysis^1,2^ were retrieved from (a) the Cancer Cell Line Encyclopedia (CCLE)^3^, (b) the Genentech Cell Line Screening Initiative (sGSI)^4^, (c) a repository of selected cell lines assembled at Columbia University, and (d) a repertoire of triple-negative breast cancer cell line profiles generated at OHSU^5^ and made available by Dr. Joe Gray. OncoMatch was designed to assess the fidelity of a model to a tumor based on the consensus overlap between their most differentially active proteins (candidate MRs) and has been shown to effectively identify cell lines representing high-fidelity cognate models of *in vivo* human tumors that recapitulate their response to small molecule inhibitors. OncoMatch was used to identify high-fidelity models for 15 human malignancies—including models that recapitulate the MR activity of molecularly distinct subtypes—as well as for aggressive tumors from patients enrolled in the N of 1 study at Columbia University^6^, see below. Extending beyond TCGA is important because this resource is limited to primary, chemonaïve tumor samples, whose MR dependencies are substantially different from those of aggressive, treated, metastatic tumors, as represented in the N of 1 study. Use of "N of 1" samples is also relevant because PanACEA-based drug predictions were extensively validated in patient-derived xenografts (PDX) models^6^. Additional criteria, such as growth kinetics in culture, cell line availability, and contamination with mycobacterium were used to prioritize among available high-fidelity models.

*1. Anaplastic Meningioma:* This malignancy is not represented in TCGA or CCLE. We thus selected two public datasets—GSE183653 (n = 185 samples) and GSE101638 (n = 42 samples)^7,8^. Analysis of three meningioma cell lines available at Columbia identified all of them as equivalent high-fidelity models. Among these, IOMM was selected as the highest-fidelity model for the Columbia N of 1 study patient^6^ (**Figure S15A**).

*2. Bladder Adenocarcinoma (BLCA):* TCCSUP was identified as the highest-fidelity model for the bladder cancer patient in the Columbia N of 1 study^6^.

*3. Breast Adenocarcinoma (triple-negative subtype) (TNBC):* HCC1143 and BT20 were identified as complementary, high-fidelity models for most TNBC samples in the TCGA and Seoul National University Hospital (GSE58135)^9^ TNBC cohorts (**Figure S15B**).

*4. Colorectal Adenocarcinoma (COAD)*: LoVo was identified as a high-fidelity model for most samples in the TCGA-COAD and TCGA-READ cohorts ([https://www.cancer.gov/tcga](https://www.cancer.gov/ccg/research/genome-sequencing/tcga)) (**Figure S7C**).

*5. Gastroenteropancreatic Neuroendocrine Tumors (metastatic) (GEP-NET):* HSTS and KRJ1 were previously identified as high-fidelity models for a large cohort of metastatic GEP-NET tumor samples^2^.

*6. Gastrointestinal Stromal Tumors (GIST):* GIST is not included in TCGA and CCLE. As a result, GIST430 and GISTT1 were identified as optimal matches to KIT-inhibitor sensitive and resistant GIST tumors based on OncoMatch analysis of the GIST patient in the Columbia N of 1 study^6^.

*7. Glioblastoma (GBM):* The HF2597 neurosphere and the U87 cell line were identified as high-fidelity models for the proneural and mesenchymal subtype of GBM, respectively, as defined by prior TCGA studies^10^ (**Figure S16A**).

*8. Lung Adenocarcinoma (LUAD):* H1793 was identified as a high-fidelity model for the Columbia N of 1 study patient^6^ (**Figure S16B**).

*9. Neuroblastoma (NBL):* NLF was previously identified as a high-fidelity model for high-risk, MYCN-amplified tumors in the Therapeutically Applicable Research to Generate Effective Treatments (TARGET) and Dutch National Research Council (NRC) neuroblastoma cohorts^11^.

*10. Pediatric Osteosarcoma (OST):* MSKOST and SAOS2 were identified as complementary, high-fidelity models for the TARGET osteosarcoma cohort (**Figure S17A**).

*11. Ovarian Serous Adenocarcinoma:* EFO21 was identified as a high-fidelity model for two metastatic patients in the Columbia N of 1 study^6^ (**Figure S17B**).

*12. Pancreatic Ductal Adenocarcinoma (PDAC):* PANC1 and ASPC1 were identified as complementary, high-fidelity models providing optimal joint coverage of basal and classical PDAC patients, respectively, in several cohorts, including TCGA-PDAC, ICGC-PDAC, E-MTAB-6830-PDAC, ORGANOID-PANCREATIC-PAAD and CPTAC-3-PAAD^12-14^ (**Figure S17C**).

*13. Prostate Adenocarcinoma (PRAD):* LNCAP and DU145 were previously identified as complementary, high-fidelity models for primary and metastatic, castration resistant prostate cancer (mCRPC)^15^ in the TCGA the Stand Up To Cancer-Prostate Cancer Foundation Cohort (SU2C)^16^, respectively.

*14. Pediatric Sarcoma:* SKNEP and MSKRMS were identified as complementary, high-fidelity models providing optimal coverage of 82 pediatric sarcoma patients’ samples collected at Memorial Sloan Kettering Cancer Center (**Figure S18A**).

*15. Stomach and Esophageal Adenocarcinoma (STAD/EAD):* N87 was identified as a high-fidelity model for samples in the TCGA-ESCA and TCGA-STAD cohorts (**Figure S18B**).

**Cell line viability assessment:** Cell lines were obtained as follows:

1. MSKOST, SAOS2, MSKRMS, and SKNEP were obtained from Memorial Sloan Kettering Cancer Center
2. GIST430 and GISTT1 were obtained from Mass General Brigham
3. The HF2597 GBM neurosphere was provided by Henry Ford Health Foundation
4. KRJ1 and HSTS were gifted by Dr. Roswitha Prfagner at the University of Graz.
5. NLF was a gift from Dr. John Maris at Children’s Hospital of Philadelphia
6. EFO21 was provided by the DSMZ.
7. All additional cell lines were procured from ATCC.

MSKOST cell line was grown in SMGM-2 Media (Lonza, Catalog #: CC-3182). MSKRMS cell line was grown in MSCGM Media (Lonza, Catalog #: PT-3001). SKNEP and SAOS2 cell lines were grown in DMEM/F12 media supplemented with 10% fetal bovine serum. KRJ1 and HSTS cell lines were grown in F12/M199(+/+) (1:1) media supplemented with 10% fetal bovine serum. NLF was grown in RPMI media supplemented with 10% fetal bovine serum. GIST430 cell line was grown in IMDM media supplemented with 10% fetal bovine serum, 100nM Imatinib. GISTT1 cell line was grown in RPMI 1640 media supplemented with 10% fetal bovine serum. EFO21 cell line was grown in RPMI media supplemented with 20% fetal bovine serum and 1xMEM Non-essential Amino Acids Solution. HF2597 cell line was grown in Neurosphere media (NM) with growth factors. Neurosphere medium (NM) was prepared by adding 5 ml N2 Supplement, 1 ml BSA stock solution (250 mg/ml), 1.25 ml Gentimicin Reagent (10 mg/ml) and 2.5 ml of Antibiotic/Antimycotic to 500 ml bottle of DMEM/F12. To prepare neurosphere medium with growth factors (NMGF), 10 ml FGFb (100 mg/ml stock), 10 ml EGF (100 mg/ml stock) was added to 50 ml NM, and NMGF was made fresh before each use or kept at 4^o^C for up to 5 days. All other cell lines were cultured using prescribed conditions described on ATCC website. To determine optimal seeding for compounds titrations, 3.2 million cells were plated and viability measured using CelTiter Glo (Promega Corp.) at 24, 48, 72 and 96 hours. Briefly, 10 mL of 320,000 cells/mL cell-solution was added to column 11 of a 12-well deep-well plate. 5 mL from column 11 was then serially diluted 1:1 from column 11 through column 2. The Hamilton MicroLab automated liquid handling system’s Cell Line Optimization protocol was next used to split each of the 12-well plates across four 384-well plates for incubation. 384-well plates were stored in the incubator and at 24, 48, 72 and 96 hours 1 plate was removed and allowed to sit for 15 minutes at room temperature. 25 μL of Cell Titer Glo was added to each well and shaken at 800 rpm for 5 min. Finally, luminescence was read using a luminometer.

**Compound titration:** To determine the 48h EC_20_ of each drug in each cell line, cells were plated into 96-well, white tissue-culture plates, in 100 µL total volume, and incubated at 37°C. After 16 hours plates were removed from the incubator and compounds were transferred into assay wells (1 μL) in triplicate. Plates were then returned to the incubator. After 48 hours the assay plates were removed from the incubator and allowed to cool to room temperature prior to the addition of 100 μL of CellTiter-Glo (Promega Inc.) per well. The plates were then mechanically shaken for 5 minutes prior to readout on the EnVision Multi- Label Reader (Perkin Elmer Inc.), using the enhanced luminescence module. Relative cell viability was computed using matched DMSO control wells as reference. EC_20_ was estimated by fitting a four-parameter sigmoid model to the individual readouts.

**Perturbed RNA-Seq profiling using PLATE-Seq:** Using the previously described plating and perturbation procedure we perturbed each cell-line with each drug at its 48h EC_20_ value (as measured above) or its C_Max_ concentration as an upper bound—defined as the maximum plasma concentration after the administration of the drug at the maximum tolerated dose in patients, (whenever available from published pharmacokinetic studies). The objective was to optimize the clinical translation potential of the PanACEA database. A few exceptions included drugs that did not demonstrate any significant effect on cell viability at concentrations ≤ 10 μM; these drugs were titrated at the 10 μM concentration. The mRNA from perturbed cells was isolated and profiled using the standard PLATE-Seq technology^17^ protocol, at 24h after perturbation.

**RNA-Seq data processing and quality control:** RNA-Seq reads derived from each well were mapped to the human reference genome assembly 38 using STAR aligner^18^. Counts files were aggregated across all 96 wells per plate and quality control (QC) reports were generated for each plate. Each plate’s QC report combined mapping statistics (STAR log files), library complexity analysis, and analysis of well-specific cross contamination using a spike-in RNA library (ERCC92 Ambion) added on alternating columns of the 96 well plate, as discussed in the PLATE-Seq protocol.

Individual plate counts files were then combined, normalized, and transformed. Specifically, First, individual counts files were merged across genes and ERCC2 spike-in counts were removed, thus yielding raw counts file for each cell-line experiment. Second, raw counts were quantile normalized and variance stabilized, based on the negative binomial distribution with the DESeq2^19^ R package (Bioconductor). To account for plate-based batch effects (which are common with drug-perturbed transcriptomic data) normalized expression was batch-corrected using ComBat^20^. Then, differential gene expression was assessed by Student’s T test analysis of drug vs. vehicle control-treated wells, pooled across all cell-line plates. Individual differential gene expression signatures were then transformed into protein activity signatures using the VIPER algorithm^21^, using tissue-of-origin-matched, context-specific regulatory network generated by the ARACNE algorithm^22^. Quality Control reports were generated across cell-lines and include multi-dimensional scaling to track the global effects of each transformation step across on each set of plates.

**ARACNE network generation:** We used the ARACNe algorithm to generate context-specific regulatory networks, using datasets comprising ≥ 100 RNA-Seq profiles of human cancer tissues from TCGA. For meningioma (GSE212377) and neuroendocrine tumors (GSE98894), we used datasets generated at Columbia University, see Table S2 for details on matched cell lines, interactomes, and cohorts used for these analyses.

TCGA RNA-Seq raw counts data were downloaded from NCI Genomics Data Commons^23^ and subsequently normalized and variance stabilized by DESeq2 R package. The ARACNe analysis included a set of 1,877 transcription factors, 677 transcriptional cofactors, and 3,895 genes encoding for signal transduction proteins as defined in the Gene Ontology (GO)^24^. Specifically, for transcription factors we selected genes in “DNA-binding transcription factor activity”, “DNA binding”, “transcription regulator activity”, “regulation of transcription” and “DNA-templated” categories (GO:0003700, 0003677, 0030528, 0003677 and 0045449, respectively). For transcriptional cofactors we selected genes in “transcription coregulator activity” plays a role in “regulating transcription” or “regulation of transcription” categories (GO:0003712, 0030528 and 0045449, respectively). Signal transduction genes were selected from those annotated as “Signal Transduction” (GO: 0007165) in the “Biological Process” section and as “intracellular” (GO: 0005622) or “plasma membrane” (GO: 0005886) in the “Cellular Compartment” section.

ARACNe was run with 100 bootstrap iterations with default parameters, including data processing inequality (DPI) set as 0 and mutual information (MI) threshold set to P = 10^−8^. The mode of regulation was computed based on the correlation between regulator and target gene expression as described in previous publications^21^.

**VIPER analysis improves data reproducibility:** For each cell line/drug replicate pair, we measured the enrichment of genes that were significantly differentially expressed in drug vs. vehicle control-treated cells in one replicate (FDR < 0.1, Benjamini-Hochberg corrected) in genes that were differentially expressed in drug vs. vehicle control-treated cells in the other replicate, as assessed by Gene Set Enrichment Analysis (GSEA)^25^. Then we performed the same analysis using VIPER-inferred protein activity rather than gene expression, using the top 50 most differentially activated and 50 most inactivated proteins by a drug. The results of this analysis, for each cell line, are summarized as violin plots, representing the distribution across all drug perturbations (**Figure S2 A-W**).

**Drug Similarity Score (DSS_AB_) calculation:** We assume that a drug’s MoA is recapitulated by the proteins whose differential activity (both positive and negative) in drug vs. vehicle control-treated cells is statistically significant. As such, given two drugs, Rx_A_ and Rx_B_, we assess their Drug Similarity Score in a specific cell line *L* (${DSS}_{A,B}^{L}$) as follows. Let’s define ${DSS}_{A\to B}^{L}$ as the Normalized Enrichment Score (NES) of the 50 most differentially activated and 50 most differentially inhibited proteins by Rx_A_ in proteins differentially activated and inhibited by Rx_B_ in the cell line $L$, as assessed by the aREA algorithm^21^. The latter represents an analytical extension of the GSEA algorithm^25^ that has been extensively validated in multiple previous studies and shown to outperform both Fisher exact test and GSEA^21^. Then we compute ${DSS}_{A,B}^{L}$ as the average of ${DSS}_{A\to B}^{L}$ and ${DSS}_{B\to A}^{L}$. The reason for selecting the 100 most differentially active proteins (candidate MRs) is two-fold: (a) using a fixed number of candidate MRs allows direct comparison of the NES between different drug pairs and (b) we have shown that the top 50+50 MRs recapitulate the proteins that control the transcriptional state of a cell^26^. In conclusion, to compute $DSS_{A,B}$ for two drugs (Rx_A_ and Rx_B_), we first define $DSS_{A\to B}$ as the enrichment of the most differentially active proteins in a cell line following treatment with Rx_A_ in proteins differentially active following treatment with Rx_B_, both compared to vehicle control. The symmetric Drug Similarity Score metric ($DSS_{A,B}$) is then the average of $DSS_{A\to B}$ and $DSS_{B\to A}$.

**Cell line MoA similarity score (CMSS_L1L2_) calculation:** Consider a specific drug (Rx) used to perturb a subset of the PanACEA cell lines ${(L}_{1},L_{2},\ldots,L_{N})$. First, for each cell line, we generate an Rx-specific protein activity vector, representing the drug-mediated differential activity of all TFs, co-TFs, and signaling proteins as assessed by VIPER in drug vs. vehicle control-treated cells. We then calculate the pairwise MoA similarity score for each ${(L}_{i},L_{j})$ pair, using the aREA gene set enrichment analysis algorithm. Specifically, the Cell line-specific MoA similarity score (${CMSS}_{i\to j}$) for a pair of cell lines ${(L}_{i},L_{j})$ is computed as the Normalized Enrichment Score (NES) of the 50 most differentially activated and 50 most differentially inhibited proteins in Rx treated $L_{i}$ cells in proteins differentially activated and inhibited by the same drug in $L_{j}$ cells, respectively. Then the Cell Line-specific Similarity Score ${CMSS}_{i,j}$ of a drug in two cell lines ${(L}_{i},L_{j})$ is defined as the average of ${CMSS}_{i\to j}$ and ${CMSS}_{j\to i}$. The resulting ${CMSS}_{i,j}$ matrix is then used to identify cell line clusters presenting similar intra-cluster MoA for a specific drug, as discussed in the next section. The “viperSimilarity” and “scale” functions from “viper” package^21^ are used to perform these analyses. In conclusion, to compute $CMSS_{C_{1},C_{2}}$ as a metric to assess the eMoA similarity of the same drug in two different cell lines, we first define $CMSS_{C_{1}\to C_{2}}$ as the enrichment of the most differentially active proteins following $C_{1}$ cells treatment in proteins differentially active following $C_{2}$ cells treatment, both compared to vehicle control. The symmetric Cell Line MoA Similarity Score metric ($CMSS_{C_{1},C_{2}}$) is then defined as the average of $CMSS_{C_{1}\to C_{2}}$ and $CMSS_{C_{2}\to C_{1}}$.

**Cell line clustering:** We used “cluster.hierarchy.dendrogram” function from “Scipy” package^27^ to perform cell line clustering analysis. The input to this analysis is the ${CMSS}_{i,j}$ matrix, generated as discussed in the previous section, and a dendrogram threshold $D_{T}=1$. Specifically, all cell line pairs with pairwise distances smaller than $D_{T}$ are clustered together and assigned the same color in the dendrogram. Cell lines forming no pairs with distances smaller than the threshold are included into the “C0” group, and will not be considered to effectively cluster with any other cell line. Only drugs that were used to perturb ≥ 2 cell lines are considered in this analysis.

**Assessing global Drug MoA similarity:** Based on the previous analysis, each drug (Rx) will be associated with a set of cell line clusters $C_{Rx}={(C}_{1},C_{2},\ldots,C_{N})$, with each cluster comprising a subset of cell lines with conserved MoA. To assess a global MoA similarity between two drugs ${Rx}_{i}$and ${Rx}_{j}$, across all perturbed cell lines, we need to assess whether their MoA is conserved in the subsets of cell lines where each drug presents individually with a conserved MoA. To achieve this goal, we first compute the intersection of all possible cluster pairs $C_{i,j}=C_{i}\cap C_{j}$), representing subsets of cell lines with conserved MoA for each drug. We then remove all intersections with Jaccard index < 1/3—*i.e.*, with overlap lower than 50% or with fewer than 2 cell lines.

For instance, consider drug ${Rx}_{A}$ with conserved MoA in clusters $C_{A,1}=(L_{1,}L_{3,}L_{5,}L_{10})$ and $C_{A,2}=\left( L_{2,}L_{4,}L_{6,}L_{7,}L_{8,}L_{9} \right)$ and drug ${Rx}_{B}$ with conserved MoA in clusters $C_{B,1}=(L_{1,}L_{5,}L_{9,}L_{10})$ and $C_{B,2}=\left( L_{7,}L_{8} \right)$. Then:

$C_{A1,B1}=C_{A,1}\cap C_{B,1}=(L_{1,}L_{5,}L_{10})$,

$C_{A1,B2}=C_{A,1}\cap C_{B,2}=\emptyset$ ,

$C_{A2,B1}=C_{A,2}\cap C_{B,1}=(L_{9})$ and

$C_{A2,B2}=C_{A,2}\cap C_{B,2}=(L_{7,}L_{8})$.

In this case the two drugs will be connected by two edges in the MoA similarity network, computed by Stouffer’s integration of their DSS on cell lines (L_1_, L_5_, L_10_) and cell lines (L_7_, L_8_), respectively. Clusters $C_{A1,B2}$ and $C_{A2,B2}$ will be ignored as they do not satisfy the Jaccard index constraint.

A final, integrated graph can be generated by selecting the most statistically significant of the subcluster-integrated DSS values, leading to a more compact network where drug pairs with conserved MoA in a cell line cluster are connected by a single edge.

**Pleiotropic correction:** To reduce the contribution to the DSS by proteins that are pleiotropically affected by many drugs (*e.g.*, those involved in cell stress or cell death pathways), we assign a cell line specific weight $w\in[0,1]$ to each protein, inversely proportional to the absolute value of the median NES for its differential activity, as assessed across all drugs. Thus, proteins that are consistently activated or inactivated by many drugs in a VIPER matrix will have lower weight and thus lower contribution to their MoA similarity. More specifically, we performed a linear fitting between the rank and the median of the NES of each protein across all drug perturbations. The slope of this line will track the average of the NES distribution as a function of the rank. As a result, the difference between a protein NES and the value predicted by the linear fit for a given rank, the fit's residual, $\Delta_{NES}(\mathrm{rank})$, provides a quantitative assessment of the protein’s systematic activation or inactivation across all drugs. We then used a sigmoidal function to transform the absolute value of the residuals |$\Delta_{NES}$| to the [0, 1] range, with slope s = -20 and inflection t = 0.5.

**MoA modularity analysis:** We assembled the MoA similarity network by creating edges between drugs based on the statistical significance of their global DSS, as described in the previous sections. We then performed modularity analysis to identify functional modules presenting strong intra-module MoA similarity using the Cluster ONE algorithm^28^. The rationale for using this algorithm is two-fold: First, based on the analysis protein-protein interaction network, ClusterONE was shown to outperform other graph theory-based algorithms in identifying protein complexes. This was assessed by benchmarks that used gold standard protein complexes dataset according to experimental studies^29,30^. Second, the algorithm supports fuzzy clustering, such that a node may be assigned to one or more clusters. The latter is critical to assess drug polypharmacology, a key goal of this study. Most graph theory-based clustering algorithms, including ClusterONE, require networks with edges representing interaction probabilities in the [0, 1] range. Since all the edges in the MoA networks are statistically significant, they could not be assigned a probability < 0.5. As a result, we used linear interpolation to map DSS scores into the [0.5, 1] range. This was accomplished by setting the maximum DSS to 1 and the minimum DSS to 0.5.

The research community focusing on protein complexes has generated remarkable efforts aimed at the manual curation of “gold standard” protein complex repertoires for mammalian organisms, such as CORUM^31^, which is frequently used for algorithm benchmarking and parameters optimization. The field of pharmacology lacks such systematic gold standard datasets. For instance, even though DrugBank^32^ provides functional annotation for many small molecule compounds, many of the ones profiled in the PanACEA study are either missing or their MoA is undefined or incomplete. Thus, rather than relying only on this resource, we performed additional manual curation of PanACEA drugs by searching for any available literature information (including DrugBank), thus improving assessment of an experimental “gold standard” for drug MoA set for performance analysis. We subsequently optimized the ClusterONE hyperparameters and the DSS cut-off by benchmarking against this “gold standard” set, based on state-of-art cluster level evaluation metrics adopted from the network biology community^28,33^. Similar to previous studies on protein complex prediction and evaluation^28,34,35^, we defined a composite score as the sum of the overlap ratio, maximal matching ratio, and accuracy score—see below for a definition of the individual metrics. Thus, the higher the composite score, the higher the quality of the drug module map analysis.

*Overlap ratio (O):* this is a simple measurement representing the precision of predicted drug modules when compared to drugs with shared MoA in the gold standard dataset. Specifically, we first compute a score $O\left( A,B \right)$ to assess the overlap between a predicted drug module (a) and a conserved MoA module (B) in the curated “gold standard” dataset as follows:

$$O\left( A,B \right)= \frac{\left| A\cap B \right|^{2}}{\left| A \right|*\left| B \right|}$$

Here, |A| and |B| represent for the number of drugs in A and B, respectively. Consistent with previous publication, we consider two modules as matching if the overlap score is greater than 0.25. This is because at least half of the drugs in two same-size modules would have to be identical for this cutoff to be reached. The overlap ratio is thus the fraction of predicted modules matched to at least one module in the curated “gold standard” drug module map.

*Accuracy score (Acc):* this metric represents the geometric mean of sensitivity (Sn) and positive predictive value (PPV), indicating the tradeoff between the two metrics^33^:

$$Acc=\sqrt{Sn*PPV}$$

In the following section, we consider a set **A** = (*A_1_, A_2_, …, A_m_*) of predicted drug modules which are compared to a set **B** = (*B_1_, B_2_, …,* B*_n_*) of gold standard modules, with *t_i,j_* denoting the number of drugs found in both module A*_i_* and *B_j_*. The sensitivity (Sn) is the fractions of drugs in the predicted module map that are also included in the reference module set.

$$Sn=\frac{\sum_{i=1}^{n} {max}_{j=1}^{m} t_{i,j}}{\sum_{i=1}^{n} {|B}_{i}|}$$

*Positive predictive value (PPV):* This metric reflects how the specificity and completeness of the match between predicted drug modules and gold standard modules. A value of 1 identifies predicted drug modules that match one and only one gold standard set module. A low PPV score indicates that the predicted drug module set has poor and non-specific overlap with the gold standard module set.

$$PPV=\frac{\sum_{j=1}^{m} {max}_{i=1}^{n} t_{i,j}}{\sum_{j=1}^{m} \sum_{i=1}^{n} t_{i,j}}$$

*Maximum matching ratio (MMR):* Finally, this metric was introduced by Nepusz et al.^28^ to overcome the limitations of PPV. Specifically, if predicted modules overlap with multiple gold standard modules, then the PPV would be low. But module overlap is important as it may pinpoint critical polypharmacology. The MMR score represents an attempt to address this issue:

$$MMR=\frac{\sum_{i=1}^{n} {max}_{i=1}^{m}O\left( n_{i},m_{j} \right)}{|{max}_{i=1}^{n}O\left( n_{i},m \right)>0|}$$

We thus used the sum of overlap ratio, accuracy score and maximum matching ratio as the composite score to evaluate the hyperparameters combination and DSS threshold cutoff. The hyperparameters searching space for clustering algorithm include (a) the density score, which was explored in the [0.3, 0.91] range, in 0.01 increments, (b) the overlap score, which was explored in the [0.1, 0.75] range, in 0.025 increments, and (c) the percentage cutoff for DSS which was explored in the [5%, 20%] range, in 0.5% increments. The hyperparameter combination that gives the highest composite score was: overlap score = 0.375, density score = 0.45 and percentage cutoff for DSS = 10%.

**Module Consistency Assessment:** Three criteria were used to ensure the internal consistency of predicted drug modules. First, we computed a z-score representing the confidence assignment of an individual drug ${Rx}_{i}$ to a module $M$ comprising $N$ drugs in total (*i.e.*, the specificity of a drug’s assignment to a module). A drug modularity score ($DMS\left( Rx_{i},M \right)$) was computed by integrating the individual $DSS_{i},_{j}$ z-scores (NES values) between $Rx_{i}$ and all other $Rx_{j=1,\ldots,N-1}$ drugs in the module, using Stouffer’s method. To assess the statistical significance of the $DMS\left( Rx_{i},M \right)$ of a specific module drug, we generated two non-parametric null hypothesis models. We also computed a global modularity score $(DMS\left( M \right))$ for each drug module, $M$, by integrating the $DMS\left( Rx_{i},M \right)$ of each individual drug in the module, using Stouffer’s method.

*1.* ***Drug Specificity Test:*** *Probability* $DMS^{NH}(Rx_{i}.N)$ *that the MoA of* $Rx_{i}$ *is similar to that of N – 1 random drugs*: For this purpose, we computed the ${DSS}_{i,j=1\ldots N-1}^{NH}$ representing the null hypothesis MoA similarity between $Rx_{i}$ and $N-1$ randomly selected drugs $Rx_{j}\notin M$ (i.e., not in the M module), producing an integrated $DSM^{NH}\left( Rx_{i},N \right)$, by Stouffer’s integration, whose value depends on both $Rx_{i}$ and $N$. This process is repeated 10,0000 times to generate a DMS probability density function (PDF) by fitting a Gaussian PDF. The PDF is used to assess the statistical significance of each $DMS(Rx_{i},M)$ score. A different null hypothesis PDF was generated for every $(Rx_{i},M)$ combination observed in the predicted module set. Drugs that failed to generate scores larger than the corresponding statistical significance threshold ($p\leq0.05$) were considered to be poor module matches.

*2.* ***Module Specificity Test:*** *Probability* $DMS^{NM}(Rx.M)$ *that a randomly selected drug (*$Rx_{j}$*) presents MoA similarity to the* $N$ *drugs in module* $M$: For this purpose, we first computed the ${DSS}_{i,j}$ between each ${Rx}_{j}\notin M$ (*i.e.* drugs not in the M module) and each drug ${Rx}_{i}\in M$ (*i.e.*, in the module). Then we integrated the $N$ individual ${DSS}_{i=1,\ldots,N,j}$ scores, using Stouffer’s method, to generate a null hypothesis modularity score $DMS(Rx_{j})$, representing the likelihood that a drug not in the module may belong to it. Finally, we generated a PDF by fitting a two-component Gaussian mixture model to all the $DMS(Rx_{j})$ values and used it to assess statistical significance at a $p\leq0.05$ level.

Notably, while first null hypothesis model produces a different threshold for each drug in a module, the second one produces a single threshold for the entire module. Thus, each drug in a module must satisfy both statistical significance tests to be retained.

*3.* ***Module Significance Test:*** *Null hypothesis for the global significance of a module:* Since the modularity score $DMS\left( M \right)$ of module $M$ depends on its size, $N$, an appropriate null hypothesis model is required to assess its statistical significance. This is generated by generating null hypothesis modules ($M_{i}^{NH}$) by selecting $N$ drug at random, 10,000 times, from the entire PanACEA set and computing the associated $DSM(M_{i}^{NH})$ to generate the associated PDF and statistical significance threshold. A potential limitation of this methodology is that same size modules will have virtually identical significance threshold, independent of drug composition.

Overall, we noticed that the last test is satisfied by virtually all predicted modules, with only one exception, which was excluded from the final module map. Any drugs that did not satisfy the first two statistical significance tests were removed from the corresponding module. For drugs that satisfied one test but not the other, we carefully checked the drug information and literature to decide whether the drug was assigned to the correct module. We identified 7 drugs meeting these criteria; these are shown using a different color in Table S6. As such, the readers should consider these results with caution.

**Mechanisms affecting drug eMoA analysis:** We downloaded the mutated gene data from DepMap portal for all overlapped cell lines in PANACEA and DepMap (n = 14). Under the same drug, for each cell line cluster and mutation combination, we constructed a 2 by 2 contingency table and performed Fisher’s Exact test to check if any mutation is statistically significantly enriched in a particular cell line cluster. To ensure the accuracy of statistical test, we only consider mutations that are observed in at least 3 cell lines and all the derived p values are corrected by Benjamini-Hochberg (BH) procedure.

For perturbation using drugs from the same chemotroxic drug group, we first computed symmetric Cell Line MoA Similarity Score metric ($CMSS_{C_{1},C_{2}}$) for any pair of TP53 mutated cell lines, and then computed ($CMSS_{C_{1},C_{2}}$) for all other pairs of cell lines (TP53 mutated vs. TP53 wild type and TP53 wild type vs. TP53 wild type). And then we performed Mann-Whitney U test to test if the two groups of $CMSS_{C_{1},C_{2}}$ scores are statistically different, and all the derived p values are corrected by Benjamini-Hochberg (BH) procedure. The same analysis is subsequently applied to all mutations.

**PrePCI, Predicting Protein-Compound Interactions:** The PrePCI algorithm^36^ extends the PrePPI^37^ algorithm to identify candidate protein/compound interactions by evaluating: a) the structural similarity of the query protein to that in a protein/compound complex found in the PDB^38^ database and, b) the chemical similarity , measured by the Tanimoto coefficient of the query compound to the one found in. the PDB complex. To elucidate potential mechanisms supporting drug polypharmacology or novel drug MoA, we queried the PrePCI database^36^ of about 5 billion predictions of protein/compound interactions for outlier compounds in Modules 1-6. The data base query was initiated using the PubChem identifiers (PMID 39558165) of each compound, as described^36^. False Positive Rate (FPR) ranges were derived by evaluating the PrePCI predictions on a PubChem-derived gold standard set. PrePCI predictions can be retrieved from <https://honiglab.c2b2.columbia.edu/prepci.html>.

For the analysis of uncharacterized compounds in Module M1, which was almost exclusively comprised of PI3K/AKT/MTOR inhibitors, potential relevant binding proteins were identified as those included in the MSigDB^39^ gene sets named: (a) PID_PI3KCI_AKT_PATHWAY, (b) REACTOME_PI3K_AKT_SIGNALING_IN_CANCER, (c) REACTOME PI3K_AKT_ACTIVATION, (d) KEGG_MTOR_SIGNALING_PATHWAY, (e) BIOCARTA_MTOR_PATHWAY, and (f) REACTOME MTOR_SIGNALLING.

For the analysis of uncharacterized compounds in Module M4, which was almost exclusively comprised of MAPK inhibitors, candidate binding proteins were identified as those in the MSigDB gene set GOBP_MAPK_CASCADE.

PrePCI was used as a supporting approach to elucidate physical binding proteins that contribute to the predicted drug polypharmacologies. We also compared PrePCI results with thermal shift assay results. In this case, we only consider proteins detected from thermal shift assay with an adjusted p-value <= 0.01 and PrePCI predictions with a high structural and binding site score (LT-scanner >= 0.3) and a moderate chemical similarity (Tanimoto Coefficient > 0.5). For both KW2449 and Az12419304, PrePCI targets are highly enriched for functional annotation terms consistent with the known MoA of their modules. For all three drugs, PrePCI targets have substantial overlap with the thermal shift proteins (24% to 67%).

For KW2449 (Module 1, PI3K/AKT/mTOR inhibitors): Consistent with the known MoA of the module, there are over-representation of the PrePCI targets results in "PI3K-Akt signaling pathway" (KEGG) with an adjusted p-value = 7.7×10^-14^ by GSEA. For the 90 thermal shift proteins with an adjusted p-value <= 0.01, PrePCI makes predictions for 60 (67%).

For Az12419304 (Module 3, MEK inhibitors): Consistent with the known MoA of the module, over-representation of the PrePCI targets results in "MAPK signaling pathway" (KEGG) with an adjusted p-value = 6.8×10^-42^ and "MAPK cascade" (GO:BP) with an adjusted p-value = 1.4×10^-31^.  For the 45 thermal shift proteins with an adjusted p-value <= 0.01, PrePCI makes predictions for 12 (27%).

**Drug-protein docking:** Molecular docking studies were performed using the 2024-2 release of the Schrödinger suite of tools. The initial poses for BRAF and RPS6KB1 in complex with their respective template ligands were obtained from the PrePCI database^40^, whereas the GPI complex structure was obtained from the PDB file 7WLD. SMILES strings for each of the predicted small molecules of interest were obtained from the PubChem Database. Protein structures were prepared using Maestro’s Protein Preparation Workflow tool with default settings^41^. Ligand structures were prepared using the LigPrep tool; chirality was inferred from 3D coordinates for template ligands while chiralities specified in the query ligand SMILES strings were maintained, otherwise ligands were prepared with default settings. Receptor grids were generated around the ligand binding sites using the Receptor Grid Generation tool; all neighboring groups were set as rotatable for BRAF and RPS6KB1 while no neighboring residues were set as rotatable for the GPI complex due to the significant number of adjacent groups. Docking for each protein-ligand pair was then performed using the Ligand Docking tool with the SP forcefield and otherwise default parameters^42^. For all three complexes, Induced Fit Docking with Molecular Dynamics (IFD-MD) was performed using the docked pose with the best docking score as the starting pose to be optimized^43^.

The Schrodinger software provides numerical estimates of binding affinities, and these are reported in the main text. Affinity scores less than ~-10.0 kcal/mol are often associated with positive predictions.

**Drug MoA and polypharmacology validation:** The following sections descried the details pertinent to the experimental validation of predicted drug polypharmacology or MoA.

**Cell culture:** All mammalian cells were cultured following ATCC recommended protocols at 37 °C and 5% CO_2_, in medium supplemented with 10% fetal bovine serum (HI-FBS, Thermo Fisher Scientific A5256801, Lot #2490735RP), GlutaMAX supplement (Thermo fisher Scientific 35050061) and 1% Penicillin-Streptomycin (Thermo Fisher Scientific 15140163). N87 cells were cultured in RPMI-1640 (ATCC 30-2001). U87 cells were cultured in Eagle's Minimum Essential Medium ([ATCC 30-2003](https://www.atcc.org/products/30-2003)). HUH-7 cells were cultured in DMEM media (Fisher Scientific 11-995-073). Saos-2 cells were cultured in McCoy’s 5A medium (Thermo Fisher Scientific 16600108) with 15% FBS and LoVo cells were cultured in F12-K media (ATCC 30-2004).

**Protein quantification:** Protein amount in cell lysate was quantified using Pierce™ BCA Protein Assay Kit (Thermo Fisher Scientific 23228) following the manufacturer’s protocol. In brief, the standard curve was generated by serial dilutions of BSA standards. Both standards and samples were added in 96-well format clear assay plate (Greiner Bio-One 655101) in duplicates and incubated with the colorimetric working solution at 37 °C for 30 min in the dark. Absorbance at 562 nm was measured and the protein concentration of sample was interpolated from the standard curve.

**Western blot:** Approximately one million cells were collected and washed twice with PBS. Cell pellet was lysed in 50 μL RIPA buffer (Thermo Fisher Scientific 89901) containing protease inhibitor cocktail (Sigma-Aldrich 11697498001) and Hult™ phosphatase inhibitor cocktail (Thermo Fisher Scientific 78420) and incubated on ice for 30 min. Lysate was centrifuged at 20,000 g for 10 min at 4 °C. The supernatant was collected, quantified, and diluted with SDS-PAGE sample loading buffer [6X] (G Biosciences 786-701) containing 120 μM DTT (Cell Signaling Technology 7722S) and boiled at 95 °C for 5 min. An equal amount of protein in the range of 25-40 μg was loaded in each lane of the NuPAGE 4-12% Bis-tris gel (Fisher Scientific WG1401 or WG1402) and transferred onto nitrocellulose membranes (Thermo Fisher Scientific IB23001) using electrophoretic semi-dry western blot transfer system. Membranes were blocked in TBS-T (TBS with 0.1% Tween20) supplemented with 5% skim milk (BD 232100) for 1 hr at room temperature and incubated with primary antibody diluted in TBS-T supplemented with 1% BSA (Sigma-Aldrich A7906) overnight at 4 °C with following dilutions:

P70 S6 kinase: (47D7) 2708S (Cell Signaling Technology), 1:1000.

Phospho p70 S6 kinase: (Thr389) (108D2) 9234S (Cell Signaling Technology), 1:1000.

p44/42 MAPK (Erk1/2): 9102S (Cell Signaling Technology), 1:1000.

Phospho-p44/42 MAPK (Erk1/2): (Thr202/Tyr204) 9101S (Cell Signaling Technology), 1:1000.

DHFR: (E6L1H) 43497S (Cell Signaling Technology), 1:1000.

Alpha tubulin: (DM1A) Sc-32293 (Santa Cruz Biotechnology), 1:2000.

GAPDH: (0411) Sc-47724 (Santa Cruz Biotechnology), 1:2000.

Membranes were washed three times in TBS-T and then incubated with fluorophore-conjugated secondary antibodies (IRDye®800CW goat anti-rabbit #926-32211 diluted at 1:4000, IRDye®680RD donkey anti-mouse #926-68072 diluted at 1:20,000; both from LI-COR) in TBS-T + 5% skim milk for 1-2 hr at room temperature. Membrane was washed three times in TBS-T and imaged on LI-COR Odyssey infrared imaging system. Images were collected and optimized contrast settings and quantified using the Image Studio software.

**Cell culture for RT-PCR and Western Blot:** The following paragraphs provide experimental details related to the validation of drugs targeting Master Regulator proteins in specific cell lines, using three complementary approaches, including qRT-PCR and Western Blots (see Figure 7). For WB and RT-PCR assays, cells were seeded into 15 cm dishes at 3.5×10^6^ cells/plate and the drugs/DMSO were added at the same time. After 24 hrs incubation with each drug at the concentration of min[EC_50_, C_max_], cells were trypsinized, washed with PBS and frozen at -80 °C until the isolation of RNA and protein. Each experiment was performed in triplicate.

**Controls selection for Western Blot and RT-PCR:** Negative control proteins were selected from the following list: ActinB, GAPDH, Tubulin Beta Class I, Tubulin Alpha 4a, Cyclophilin A, Cyclophilin B, Vinculin, HDAC1, Lamin B1, PCNA and TBP. The specific control proteins for each drug in a specific cell were identified by determining whether the specific drug treatment did not induce changes in the corresponding gene expression compared to vehicle control-treated samples (DMSO).

**Western Blot:** Approximately 90% of the total cells collected from each 15 cm dish were used for Western Blot. Briefly, cell pellets were lysed into colorless 1 x SDS-Laemmli buffer, followed by protein concentration measurements with DC Protein Assay kit II (Bio-Rad). After normalizing all the protein concentrations, equal amount of protein lysate was loaded to the wells (Bolt™ Bis-Tris Plus Mini Protein Gels, 4‐12% (Invitrogen)) of the samples which needed to be compared. Proteins were transferred to 0.2 μm Nitrocellulose membrane by using either Trans-Blot® SD Semi-Dry Transfer Cell (Bio-Rad) or by using iBlot™ 2 Gel Transfer Device (Invitrogen). After transfer, the membranes were blocked in 5% milk in PBS for 30 min followed by overnight incubation with primary antibodies with following dilutions:

C-MYC: [Y69] ab32072 (Abcam), 1:1000 in 5% milk in PBS.

CTNNB1: (D10A8) 8480 (Cell Signaling Technology) 1:1000 in 5% milk in PBS.

UHRF1: (H-8) sc-373750 (Santa Cruz Biotechnology), 1:300 in PBS.

CENPF: ab5 (Abcam), 1:500 in 5% milk in PBS.

ACTB: (C4) sc-47778 (Santa Cruz Biotechnology), 1:1000 in 5% milk in PBS.

GAPDH: (14C10) 2118 (Cell Signaling Technology) 1:1000 in 5% milk in PBS.

After overnight incubation, membranes were washed 5 x 5 min with PBS-0.1%Tween, followed by secondary antibody incubations (HRP-conjugated Donkey Anti-Rabbit IgG and Anti-mouse IgG, 1:30000 for ACTB and GAPDH, 1:10000 dilution for all other proteins) for 30 min. After the secondary antibody incubations, the membranes were again washed 5 x 5 min with PBS-0.1%Tween followed by chemiluminescent detection by SuperSignal™ West Pico PLUS Chemiluminescent Substrate (Thermo Scientific). All Western Blots were done in 3 biological replicates.

**Cell fractionation Western Blots:** The drug treatments were done similarly as with other WBs. After 24 h drug / DMSO treatments, the cells were trypsinized and collected to 15 ml falcons followed by 2 x PBS washes. The cellular fractionations were done by using NE-PER™ Nuclear and Cytoplasmic Extraction Reagents (Thermo Scientific). First, 40 ul of ice-cold CER I buffer with Protease inhibitors (cOmplete™, Mini, EDTA-free Protease Inhibitor Cocktail, Roche) was added to the cell pellets, followed by vortexing for 15 s and 10 min incubation on ice. Next 22 µL of ice-cold CER II buffer was added, followed by 5 s vortexing and 1min incubation on ice. This was followed by another brief 5 s vortexing and centrifugation for 5 min at 16,000 × g at 4 °C. At this point the cytoplasmic extract was collected (80% of the total supernatant). The remaining 20% of supernatant was carefully discarded. To minimize cytoplasmic protein carry-over to nuclear extract, 1 × PBS (ice-cold) wash was performed, followed with centrifugation of 600 x g for 3 min at 4 °C. The remaining pellet was suspended with ice-cold NER with protease inhibitors (cOmplete™, Mini, EDTA-free Protease Inhibitor Cocktail, Roche) followed by 15 s vortex and this 15 s vortexing was continued every 10 min for total of 40 min. The whole-time samples were stored on ice. Finally, the sample was centrifuged at 16,000 × g for 10 min and supernatant (nuclear extract) was collected. Samples were stored in -80 °C until protein measurement with Dilution-Free Rapid Gold BCA Protein Assay Kit (Pierce) and WBs.

**EC-17 Uptake Assay with Mgcd265 Treatment:** Lovo cells were cultured according to manufactures recommendations: F-12K Medium with 10% FBS and pen/strep. A total of 7.3 × 10⁶ LoVo cells were seeded in 15-cm culture dishes in complete medium containing either 1 μM MGCD265 (Glesatinib; corresponding to IC₂₀) in 0.39% DMSO or 0.39% DMSO alone as vehicle control. After 24 h of incubation at 37 °C, cells were detached using Accutase, pelleted by centrifugation, and resuspended in fresh medium containing the same concentrations of MGCD265 and DMSO, supplemented with 2 μM EC-17 (disodium salt; MedChemExpress). Cells were distributed at 1 × 10⁶ cells per tube and incubated at 37 °C for the following time points: 0 min (no EC-17), 30 min, 60 min, 90 min, 120 min, and 150 min. Work path for each biological replicate listed below (3 Biological replicates in total). At each time point, cells were washed three times with ice-cold DPBS and fixed in 4% paraformaldehyde for 10 min at room temperature. Fixed samples were analyzed by flow cytometry using a NovoCyte Penteon Cell Analyzer. The 0 min time point (no EC-17) was used as a gating control to define EC-17–positive populations.

**Proteome Integral Solubility Alteration Assay (PISA):** Frozen cell pellets were suspended in ice-cold lysis buffer (20 mM HEPES, 138 mM NaCl, 5 mM KCl, 2 mM CaCl_2_, 1 mM MgCl_2_, 1× complete EDTA-free protease inhibitor cocktail, final pH 7.4) and gently homogenized via 20 strokes with a tight-fitting douncer. The resulting crude lysate was centrifuged at 300 rcf for 3 minutes and supernatant was transferred to a new LoBind tube. Lysate was assayed using a Pierce BCA Protein Assay Kit and adjusted to 2.0 mg/mL with lysis buffer. Lysate was equilibrated to room temperature and mixed 1:1 with lysis buffer containing a 2 × concentration of drug or a vehicle control, yielding a 1 × lysate (final protein concentration 1 mg/mL, final drug concentration 20 µM, 4 replicates per group). The 1 × lysate was incubated at room temperature for 15 minutes then distributed across a 96-well PCR plate in 20 µL aliquots. The plate was heated in a thermal cycler for 3 minutes across a thermal gradient (48 to 64 °C), then cooled and incubated at room temperature for 5 minutes. The plate was placed on ice, then an equal volume of lysate (18 µL) was used to pool temperature points back to their original replicate samples. Samples were treated with 100 units of Benzonase on ice for 30 minutes to digest nucleic acids. The pooled samples were mixed 1:1 with lysis buffer supplemented with 0.8% NP-40 (final NP-40 concentration 0.4%), gently vortexed, and incubated on ice for 15 minutes to extract membrane proteins. Aggregates were pelleted by centrifugation at 30,000 RCF and 4 °C for 1 hour. 50 µL of supernatant was transferred to new tubes containing 25 µL of 3 × sample preparation buffer (300 mM HEPES pH 8.5, 6% SDS, 30 mM TCEP, 120 mM chloroacetamide), vortexed, heated to 95 °C for 15 minutes, then cooled to room temperature. Samples were processed using protein aggregation capture (PAC).

**PISA sample processing:** For PAC-based sample processing^44^, samples were mixed with 4 µL of pre-washed carboxylate-modified magnetic bead slurry (SergiLabs^45^) and proteins precipitated using acetonitrile (70% v/v). Samples were incubated on a ThermoMixer (24 °C, 1000 RPM) for 20 minutes, centrifuged for 2 minutes at 1,000 RCF, and beads drawn to a magnet. Liquid was removed via vacuum aspiration. Beads were washed 3 × with acetonitrile, air-dried, and resuspended in 50 µL of digestion buffer (200 mM EPPS pH 8.5, 10 mM CaCl2, 0.02 µg/µL Trypsin). Samples were digested overnight at 37 °C with vigorous shaking. Digests were reacted with ~100 µg of TMTpro reagents, quenched with hydroxylamine, pooled, and dried in a SpeedVac. The sample was resuspended in 2% acetonitrile, acidified with formic acid, and desalted using a Pierce Peptide Desalting Spin Column (Thermo Fisher). The eluate was evaporated in a SpeedVac prior to high pH reversed-phase peptide fractionation. Peptides were fractionated using high pH reversed-phase chromatography on an Agilent 1260 Infinity Capillary LC equipped with an XBridge Peptide BEH C18 column (300Å, 3.5µm, 1mm x 150mm, Waters Corporation). Peptides were resuspended in 2% acetonitrile and 0.1% NH_4_OH, injected manually, and separated using a gradient of mobile phase A (2% acetonitrile, 0.1% NH4OH) to mobile phase B (60% acetonitrile, 0.1% NH4OH) at a flow rate of 75 µL/min. 96 fractions were collected and combined into 24 pooled fractions, then evaporated in a SpeedVac, and resuspended in 2% acetonitrile, 0.1% formic acid. Peptide concentration was determined via Nanodrop.

**LC-MS/MS data collection for PISA:** Peptides (500 ng/fraction) were separated and analyzed on an Easy-nLC 1200 system connected to an Orbitrap Eclipse Tribrid mass spectrometer equipped with a FAIMS Pro Interface. Mobile phase A consisted of 2% acetonitrile, 0.1% formic acid in water and mobile phase B consisted of 80% acetonitrile, 0.1% formic acid. Peptides were first loaded onto an Acclaim PepMap C18 nano-trap column (100Å, 3*µ*m, 75*µ*m × 2cm, Thermo Scientific 164946) in mobile phase A, then resolved over a 120 minute method on an EASY-Spray column (100Å, 2*µ*m, 75*µ*m × 500mm, Thermo Scientific ES903). The flow rate was set to 250 nL/min. MS data was collected using a FAIMS-TMTpro-MS2 method. The mass spectrometer was operated in positive ion mode with a spray voltage of 2500V and a capillary temperature of 275 °C. Data dependent acquisition was performed with a cycle time of 1 second per FAIMS CV (-35, -50, -65V). MS1 scans were acquired in the Orbitrap with a resolution of 60,000, scan range of 400-1600 m/z, normalized AGC target of 100%, and maximum injection time set to automatic. Additional filters included MIPS mode set to peptide, intensity threshold of 2.5e4, allowed charge states 2-6, and a 60 second dynamic exclusion window. MS2 scans were acquired in the Orbitrap with a resolution of 50,000, HCD collision energy set to 36%, normalized AGC target of 200%, and maximum injection time set to automatic.

**PISA data processing:** Raw files were searched using the Andromeda search engine^46^ in MaxQuant software^47^ (version 2.0.3.0) against the Uniprot human complete FASTA database (downloaded February 8, 2023) and the MaxQuant contaminants database. Searches were performed using default MaxQuant settings with minor modification. Briefly, reporter ion MS2 mode was used with internal and terminal TMTpro 16plex labels corrected according to manufacturer-supplied correction values. Searches allowed for methionine oxidation and N-terminal protein acetylation as variable modifications (max. two), while cysteine carbamidomethylation was set as a fixed modification. Enzyme specificity was set to Trypsin/P with a maximum of two missed cleavages allowed. Precursor ions were searched with a mass tolerance set to 4.5 ppm and fragment ions were searched with a mass tolerance set to 20 ppm. Peptide spectrum match and protein FDRs were set at 0.01 using a target-decoy database search strategy^48^. Only unique peptides were used for protein quantification. The resulting proteinGroups.txt output file was processed in R using the Omics Notebook package^49^. Contaminants, decoys, and proteins with missing values were excluded from the analysis. Raw abundance values were log-transformed, median-normalized, and subjected to hypothesis testing using a moderated t-test followed by a Benjamini-Hochberg correction (FDR < 0.05).

**SUPPLEMENTARY FIGURES**

**Supplementary Figure S1: Comparing reproducibility of PanACEA and L1000 profiles.** We computed raw gene counts correlation (Pearson and Spearman) between technical replicates in LOVO and BT20 cell lines for both PanACEA and L1000-related profiles.

**Supplementary Figure S2: GSEA analysis comparison between gene signature and protein activity.** Each violin plot shows the enrichment analysis, assessed by GSEA analysis, either based on the most differentially expressed genes (red) or most differentially active proteins (blue), across technical replicates for each drug, aggregated across all drug perturbations. Specifically, for gene expression, we assessed the enrichment of genes that were statistically significantly differentially expressed in one replicate (FDR < 0.1, Benjamini-Hochberg corrected) in genes that were differentially expressed in another replicate for the same drug and cell line. For protein activity, we performed the same analysis but using the genes encoding for the 50 most differentially activated and inactivated proteins.

**Supplementary Figure S3: Different eMoAs for the same drug are conserved in different cell lines (clusters).** Hierarchical clustering analysis was used to identify cell clusters with highly conserved MoA for each drug perturbation, using a pairwise similarity score $DSS_{i},j\left( Rx \right)$, representing the MoA conservation of a drug ($Rx$) in the $i$-th and $j$-th cell lines. Individual clusters are identified by different colors in the dendrogram. The six drugs shown here—including crizotinib, imatinib, nilotinib, tamoxifen, topotecan and vandetanib, selected for illustration purposes—show the diversity of cell line-specific MoA clustering. Identified clusters ranged from $N=4$, for topotecan, to $N=6$ for imatinib and tamoxifen.

**Supplementary Figure S4: TP53 mutated cell lines show closer CMSS than non TP53 mutated cell lines in four different drug classes:** (A) HDAC inhibitors; (B) PI3K inhibtors (C) Chemotoxic drugs; (D) EGFR inhibitors.

**Supplementary Figure S5: Clustering results from sensitivity data:** (A) A global correlation map for overlapped drugs between PANACEA and DepMap; (B) MEK inhibitor module (M3) correlation map using sensitivity data; (C) Correlation scores distribution. Biib021 within module (M3) correlation scores are highlighted using red bars.

**Supplementary Figure S6: Examples of cell context-specific drug modules**. (A) A drug functional module consists of three DNA damage agents and two steroidogenesis inhibitors is conserved in two osteosarcoma cancer cell lines (SAOS1 and MSKRMS) and one kidney cancer cell line (SKNEP). (B) A drug functional module consists of five drugs with very distinct MoA annotations is conserved in two triple negative breast cancer cell lines (HCC1143 and BT20). The number underneath the drug name indicates the concentration (μM) used for the drug in that particular cell line.

**Supplementary Figure S7: Gene set enrichment analysis for significant hits of Mgcd265 from Proteome Integral Solubility Alteration Assay:** GSEA analysis using all proteins detected from Mgcsd265 proteome integral solubility alteration assay (adjusted p-value < 0.01) using various references. (A) KEGG. (B) Reactome. (C) WikiPathways.

**Supplementary Figure S8: Gene set enrichment analysis for significant hits of Az12419304 from PrePCI predictions:** (A) GSEA analysis against all significant proteins detected from Az12419304 proteome integral solubility alteration assay (adjusted p-value < 0.05). (B) GSEA analysis using leading edge genes from (A) against Reactome gene sets. (C) GSEA analysis using leading edge genes from (A) against WikiPathways gene sets.

**Supplementary Figure S9: Gene set enrichment analysis for significant hits of Biib021 from PrePCI predictions:** (A) GSEA analysis against all significant proteins detected from Biib021 proteome integral solubility alteration assay (adjusted p-value < 0.05). (B) GSEA analysis using leading edge genes from (A) against CORUM.

**Supplementary Figure S10: Molecular docking for Cladribine-HASP90AA1:** (A) Initial Structure of 2-chlorodideoxyadenosine (CDY) in complex with the N-domain of the ER Hsp90 chaperone GRP94, PDB ID 1QYE chain A. (B) Docked pose of CDY in complex with GRP94 obtained using Glide. (C) Glide docked pose of cladribine in complex with HSP90A11.

**Supplementary Figure S11: PanACEA-predicted inhibitors of the 37 most recurrent MR proteins in TCGA, as identified by MOMA analysis^26^.** MR inhibition heatmap showing the effect of the top 15 drugs inferred as potent MR inhibitors (*i.e.*, VS ≤ ‑3), by VIPER analysis of drug vs. vehicle control-treated cells. Rows represent the 37 MOMA-nominated MRs, while columns represent the 23 cell lines in PanACEA. For each drug/cell line pair, the heatmap shows the effect of the top 15 VIPER-inferred inhibitors (different for each heatmap), sorted left to right from the most (dark blue) to the least potent (light blue). The inactive MRs (VIPER score <= 0 in untreated PanACEA cell lines, compared to the centroid of the CCLE and Genentech cell lines repositories) are shown in light green.

**Supplementary Figure S12: Panels characterizing the three most potent inhibitors of cell line-specific MRs, across all 23 cancer cell lines.** For each MOMA-nominated MR protein and cellular context, drugs are ranked top to bottom based on the VIPER-assessed differential MR activity in drug vs. vehicle control-treated cells. The threshold for selecting potent inhibitors was set at $NES £ -3$ and only MR proteins active in the specific cell line were considered. The inactive MRs (VIPER score <= 0 in untreated PanACEA cell lines, compared to the centroid of the CCLE and Genentech cell lines repositories) are shown in light green.

**Supplementary Figure S13: Bar charts showing the 15 most potent MR inhibitors identified by VIPER analysis of drug vs. DMSO-treated cells.** Drugs are ranked top to bottom based on the number of cell lines with VIPER $NES£-3$, as assessed from drug vs. DMSO-treated cells. Only cell lines with an active ($NES\geq0$) MR were considered in this analysis and 37 MOMA-nominated MR proteins were shown here.

**Supplementary Figure S14: False negative MR inhibitor predictions: (**A) Pimasertib in IOMM cells; (B) Pimasertib in HSTS cells.

**Supplementary Figure S15 – S18:** **Cell lines selection by OncoMatch analysis.** Each heatmap shows the statistical significance of the OncoMatch analysis ($-{log}_{10}(p)$, Bonferroni corrected) assessing conservation of the top 50 most activated and inactivated MRs (Tumor Checkpoint Matching Score, TCMS) in each available cell line vs. each sample in available tumor-matched cohorts. As discussed in^6,15^, we selected a conservative statistical threshold ($p = {10}^{-5}$) to identify high-fidelity models. The color map indicates the statistical significance of all high-fidelity model/tumor matches. The Cohort Coverage Index (CCI), shown as a barplot on the right side of the heatmap, represents the fraction of tumor samples in the selected cohorts identified as high-fidelity matches for the specific cell line. Finally, the Integrative Checkpoint Matching Score (ICMS), shown next to the CCI, represents the area over the cumulative distribution for TCMS between $p<-{10}^{-5}$ and the maximum TCMS value across all cell lines. Cell lines are ranked by the sum of the CCI and ICMS metrics.

**Supplementary Table 1: Drugs and drug concentration used in each individual cell line.**

**Supplementary Table 2: Tumor-specific cohort information for the generation of ARACNe interactomes.**

**Supplementary Table 3: Cell line clustering results for all drugs.**

**Supplementary Table 4: Mutation analysis.**

**Supplementary Table 5: Drug functional network.**

**Supplementary Table 6: Predicted of drug modules and associated statistical information and manual annotations of all modules.**

**Supplementary Table 7: PrePCI prediction results.**

**Supplementary Table 8-30: Predicted protein activity (VIPER scores) for all drugs across all 23 cell lines.**
